# Trial-wise exposure to visual appetitive cues increases physiological arousal but not temporal discounting

**DOI:** 10.1101/2021.07.24.452477

**Authors:** Kilian Knauth, Jan Peters

## Abstract

Humans and many animals devalue future rewards as a function of time (temporal discounting). Increased discounting has been linked to various psychiatric conditions, including substance-use-disorders, behavioral addictions and obesity. Despite its high intra-individual stability, temporal discounting is partly under contextual control. One prominent manipulation that has been linked to increases in discounting is the exposure to highly arousing appetitive cues. However, results from trial-wise cue exposure studies appear highly mixed, and changes in physiological arousal were not adequately controlled. Here we tested the effects of appetitive (erotic), aversive and neutral visual cues on temporal discounting in thirty-five healthy male participants. The contribution of single-trial physiological arousal was assessed using comprehensive monitoring of autonomic activity (pupil size, heart rate, electrodermal activity). Physiological arousal was elevated following aversive and in particular erotic cues. In contrast to our pre-registered hypothesis, if anything, we observed *decreased* temporal discounting following erotic cue exposure. Aversive cues tended to increase decision noise. Computational modeling revealed that trial-wise arousal only accounted for minor variance over and above aversive and erotic condition effects, arguing against a general effect of physiological arousal on temporal discounting.

## 1. Introduction

Many decisions are associated with consequences that differ in temporal proximity and reward magnitude. Temporal discounting (TD), the tendency to favour smaller-but-sooner over larger-but-later rewards, is common in humans (Peters & Büchel, 2011) and many animals (Kalenscher & Pennartz, 2008). Alterations in TD are associated with a range of psychiatric conditions and problematic behaviors, including addiction, substance abuse and attention-deficit hyperactivity disorder (Amlung et al., 2019; Bickel et al., 2019; Jackson & McKillop, 2016; Wiehler & Peters, 2015). TD exhibits stability over weeks (*r* = .91; Simpson & Vuchinich, 2000), months (*r* = .77-.80; Arfer & Luhmann, 2017) and even one year (*r* = .71; Kirby, 2009), and across different testing environments (Bruder et al., 2021; Odum, 2011). Temporal discounting is therefore regarded as a trait-like characteristic (Smith & Hantula, 2008).

However, despite its trait-like stability, TD can be modulated by contextual factors (Lempert et al., 2016; Peters & Büchel, 2011). The format of time and reward information directly influences choice behavior (Lempert et al., 2016). TD is attenuated when delays are expressed in terms of the date of reward delivery (Read et al., 2005), when future rewards are paired with participant-specific episodic cues (Bromberg et al., 2017; Peters & Büchel, 2010; Rösch et al., 2021) or when reward amounts are increased (Green et al., 1997).

In the context of TD, modulatory effects of appetitive cues have also long been discussed. For example, men might discount rewards more steeply following exposure to arousing pictures of opposite-sex faces or erotica (Kim & Zauberman, 2013; Van den Bergh et al., 2008; Wilson & Daly, 2004). Such effects are often interpreted as reflecting the activation of a motivation/reward system by highly rewarding erotic stimuli, which in turn could facilitate reward approach behaviour in other domains (Van den Bergh et al., 2008). This resonates with the observation that erotic cues robustly activate reward-related brain circuits including ventral striatum and orbitofrontal cortex (Gola et al., 2016; Stark et al., 2005; Wehrum-Osinsky et al., 2014). Furthermore, primary reinforcers such as appetitive (erotic) or food cues might promote out-of-domain immediate (monetary-) reward preferences (Li, 2008; Yeomans & Brace, 2015). Participants exhibiting steeper discounting showed increased responses to positive reward feedback in ventral striatum (Hariri et al., 2006). Effects of appetitive cues in healthy participants might also share conceptual similarities with so-called *cue-reactivity* responses in addiction, reflecting increased subjective, physiological and neural responses to addiction-related cues in substance-use-disorders and behavioral addictions (Courtney et al., 2015; Starcke et al., 2018; Volkow et al., 2010).

However, there is some heterogeneity with respect to modulations of TD via affective cues. Luo and colleagues (2014) primed participants with fearful and happy faces in a trial-wise design. Fearful faces were associated with a *reduction* in TD. In contrast, Guan and colleagues (2015) showed that trial-wise presentation of aversive cues *increased* discounting, whereas Simmank and colleagues (2015) observed no effects of erotic cue exposure across lean and obese participants.

In sum, the effects of affective cues on TD are mixed. Many studies use a blocked presentation of a series of appetitive (erotic) *and*/*or* aversive stimuli to examine effects on TD (Cai et al., 2019; Kim & Zauberman, 2013). Although such designs are suitable to robustly detect cue effects on decision-making if they exist, they are ill-suited to reveal short-term behavioral or psycho-physiological changes that accompany individual decisions.

Such cue effects might in part be driven by activation of ascending catecholaminergic brainstem arousal systems, as salient stimuli, irrespective of their valence, are associated with phasic discharges of neurons in the locus coeruleus (LC), the primary noradrenergic brainstem nucleus (Bouret & Richmond, 2015; Chen & Sara, 2007; Mather et al., 2016). Phasic LC activity increases noradrenaline release across cortex, facilitating sensory stimulus processing (Howells et al., 2012; Mather et al., 2016; Moxon et al., 2007). In line with these findings, cortical responses to non-salient stimuli can be elevated when time-locked with phasic LC photoactivation (Vazey et al., 2018).

LC activity is tightly linked to pupil dilation (Aston-Jones & Cohen, 2005; Joshi et al., 2016) in humans (Murphy et al., 2014) as well as in primates (Varazzani et al., 2015). Due to the association between phasic LC activity and cortical noradrenaline release, pupil dilation is often used in conjunction with other measures such as cardiovascular (ECG) and electrodermal activity (EDA) to examine short-term changes in arousal levels (Bradley et al., 2008). Pupil responses are increased following both aversive (Kinner et al., 2017) and erotic stimuli (Finke et al., 2017) and might track ongoing task demands and choice processes (Alnæs et al., 2014; van der Wel & van Steenbergen, 2018). Finally, trial-wise arousal changes are choice-predictive during temporal discounting (Lempert et al., 2016). In sum, appetitive cues might modulate TD under some conditions (Guan et al., 2015; Luo et al., 2014) but whether such effects can be traced back to variations in short-term physiological arousal remains unclear.

Here we address this issue, expanding upon previous work in three ways. First, we applied a trial-wise design that allowed us to disentangle arousal- and valence-related effects. Second, we comprehensively monitored psycho-physiological arousal (pupil size, cardiovascular activity, electrodermal activity). Finally, we quantified arousal-related effects on individual decisions using a hierarchical Bayesian computational modeling scheme. Based on the literature (see above), we pre-registered the following hypotheses (https://osf.io/swp4m/): On the behavioural level we predicted increased TD following both erotic and aversive cues. Further, we hypothesized that arousing stimuli (irrespective of valence) would lead to increased physiological arousal (pupil dilation, skin conductance response amplitudes, heart rate deceleration). We also hypothesized that difficult trials (high decision conflict) would result in a more pronounced pupil dilation indicating high cognitive effort. Finally, we predicted that higher baseline working memory would be negatively associated with steepness of discounting behavior (Shamosh et al., 2008).

## 2. Materials and Methods

### 2.1. Participants

Thirty-five heterosexual male participants took part in the study (mean ± SD (age) = 24.3 ± 5.1; range 18-38 years). All subjects were non-smokers, fluent German speakers, reported normal or corrected-to-normal vision and had no history of neurological or psychiatric disorders. All experimental procedures were approved by the ethics committee of the German Psychological Society (DGPs), and participants provided informed written consent prior to participation in the study. Participants were recruited online and included mainly university students. A preregistered sample size of n = 29 was determined a priori via a power analysis using G*Power (Faul et al., 2007). This effect size estimate was based on previous studies investigating cue effects on temporal discounting (average effect size across three studies: Cohen’s *d* = 0.49; alpha error prob. = 0.05; power = 0.80; Kim & Zauberman, 2013; Sohn et al., 2015; Wilson & Daly, 2004).

### 2.2. Experimental set-up

Participants were seated in a shielded, dimly lit room 60 cm from a 24-inch LED screen (resolution: 1366 x 768 pixels; refresh rate: 60 Hz). The subjects placed their chin and forehead in a height-adjustable chinrest. They were instructed to minimize blinks and to focus on the screen center throughout the experiment. Stimuli were presented centrally at 600 x 600 pixels superimposed on a gray background. Stimulus presentation was implemented using Psychophysics toolbox (Version 3.0.14) for MATLAB (R2017a; MathWorks, Natick, MA).

### 2.3. Affective Cues

We screened several image databases for stimulus selection, including the International Affective Picture System (IAPS), the Nencki Affective Picture System (NAPS) as well as EmoPics (Lang, Bradley, & Cuthbert, 2008; Marchewka et al., 2014; Wessa et al., 2010). In addition, we performed a google search. We created a preliminary stimulus set consisting of 376 erotic, aversive and neutral images, which were roughly matched for image content and complexity. As all pictures displayed humans, stimuli can be considered as social cues. In a preceding pilot study, the preliminary set was rated concerning valence and arousal levels by an independent sample (n = 10). Based on those ratings we selected 288 images (96 erotic/aversive/neutral). Erotic and aversive images were comparable in their arousal levels (mean ± SD arousal = erotic: 6.98 ± 0.52; aversive: 7.04 ± 0.82, *p* = .115) but differed in terms of their valence (mean ± SD valence = erotic: 7.57 ± 0.49; aversive: 2.50 ± 0.69, *p* < .001). Neutral images differed in both dimensions (arousal: 1.59 ± 0.25; valence: 5.63 ± 0.49). Further details on the pilot study image ratings and statistics are provided in the supplement (see Figure S1).

Using MATLAB’s SHINE toolbox, images were converted to grayscale and matched according to mean intensity (mean ± SD = erotic: 0.42 ± 0.0009; neutral: 0.42 ± 0.001; aversive: 0.42 ± 0.001; *p* = .133) and contrast (mean ± SD contrast = erotic: 0.19 ± 0.01; neutral: 0.19 ± 0.001; aversive: 0.19 ± 0.01; *p =* .347). Details of physical image properties and associated analyses are depicted in the supplement (see Figure S2).

### 2.4. Data acquisition

For quantification of cue-evoked physiological responses throughout the experiment, we assessed three different measures of autonomic nervous system activity, pupil size, heart rate and skin conductance. Pupillometry data were collected using a RED-500 remote eye-tracking system (sampling frequency (SR): 500 Hz; Sensomotoric Instruments) which uses invisible infrared illumination of the retina. Heart rate and skin conductance data were acquired by Biopac systems hard- and software (SR: 2000 Hz; MP 160; Biopac systems, Inc). For cardiovascular recordings an ECG100C amplifier module with a gain of 2000, normal mode, 35 Hz low pass notch filter and 0.5 Hz/1.0 Hz high pass filter was included in the recording system. Disposable circular contact electrodes were attached according to the lead-II configuration. Isotonic paste (Biopac Gel 100) was used to ensure optimal signal transmission. For electrodermal recordings we used an EDA100c amplifier module with a gain of 5 µS/V, 10 Hz low pass filter and DC high pass filter settings. Activity was derived from the index- and middle finger of the non-dominant hand using disposable Ag/AgCl electrodes, pre-gelled with isotonic gel. Participants responses from the behavioural tasks were recorded via keyboard and mouse.

### 2.5. General procedure

Data collection was carried out on two separate testing days with an interval of 3-7 days. On day one, participants were informed about the experimental procedure and provided informed consent. They then completed a behavioral pretest, a series of questionnaires and a number of working memory tasks. On day two, participants completed the temporal discounting task with trial-wise affective picture presentation and physiological recordings.

#### 2.5.1. Behavioral pretest and subject-specific trial generation

Participants performed a short behavioral pretest (96 trials) of a temporal discounting task. On each trial, participants chose between a fixed immediate reward of 20 € (smaller-sooner, SS) and a variable delayed amount (larger-later, LL). LL-reward amounts were calculated by multiplying the SS-amount with sixteen factors (1.05, 1.055, 1.15, 1.25, 1.35, 1.45, 1.55, 1.65, 1.85, 2.05, 2.25, 2.55, 2.85, 3.05, 3.45 and 3.85), and combined with a set of six delays (2, 6, 15, 29, 62, or 118 days), yielding a total of 16 * 6 = 96 trials. Participants were instructed explicitly about the task structure and performed twelve practice trials. We then used the pretest choice data to estimate an a-priori discount-rate via Maximum-Likelihood estimation assuming a hyperbolic model (Eq. 1) and a softmax choice rule (Eq. 2) (Green & Myerson, 2004).

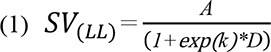

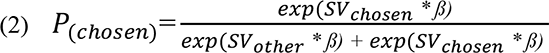

The hyperbolic model describes the decrease in the subjective value of a delayed option (LL) over time. The amount of the larger but later reward (*A*) which is delivered after a specific delay (*D* in days) is devalued by the subject’s specific discount-rate (*k*, here modeled in log-space) that weights the influence of time on the subjective value (*SV*). Higher *k*-parameter reflect an increased devaluation of the LL over time or more impulsive choice preferences. As choice preferences are affected by subject-specific noise, we used a sigmoid (softmax) function (Eq. 2) to estimate choice probabilities (Sutton & Barto, 1998). Here, *ß* scales the influence of value differences on choice probabilities. Lower values of *ß* indicate a high choice stochasticity, whereas higher values indicate that choices depend more on value differences.

We then used this pretest-based discount rate to calculate indifference points (ID-points) for each participant for three different delay vectors ([1, 7, 14, 28, 65, 90]; [1, 7, 14, 30, 55, 100]; [1, 7, 14, 32, 60, 80]). ID-points reflect the LL-amount at which a participant is expected to be indifferent between the SS- and LL-options. For every participant, the three delay-vectors were randomly assigned to the experimental conditions (erotic, aversive, neutral). The indifference amounts per delay were then used to compute participant-specific choice options. To this end, for each delay in each condition, we drew ten random samples from normal distributions centered at the respective ID-points, with standard deviation of 4, and six additional samples linearly spanning the interval between 20 and a subject-specific maximum, yielding 96 subject-specific trials per condition.

After completion of the behavioral pretest, participants underwent a short working memory test battery, including digit-(forward & backward), operation- and listening span tasks (Redick et al., 2012; van den Noort et al., 2008; Wechsler, 2008). Finally, participants completed several questionnaires on demographic, health and personality data that will be reported elsewhere.

#### 2.5.2. Temporal discounting task with affective pictures

On day two, participants performed the experimental version of the temporal discounting task including the erotic, aversive and neutral cues derived from the pilot study. Subjects were seated in a dimly lit, electrically and acoustically shielded test room with their head placed on the chinrest. After the ECG- and EDA-electrodes were attached and the eye-tracker was calibrated (9-point calibration), the discounting task was started.

Here, participants performed 288 choices between a smaller-sooner reward of 20 € immediately available and one of the subject-specific delay/LL-reward pairs generated from the pretest. The overall trial structure is outlined in Figure 1. Every trial started with the presentation of one of the 96 neutral, aversive or erotic images which were presented in the screen center for 2000ms. Then the LL-reward and the associated delay were superimposed on the image (e.g. 38 €, 14 days). After 3000ms the image and the LL-reward disappeared and the decision screen was presented. Here, participants chose between one of two symbols which corresponded to the available options (SS: circle; LL: square). The chosen option was then highlighted for another 1000ms. The intertrial-interval (ITI) was marked by a white fixation cross superimposed on a grayscale scrambled image with a randomized presentation time between 5500ms and 6000ms sampled from a uniform distribution. The scrambled image was exactly matched to the affective picture set in terms of mean pixel intensity and contrast. Experiment administration and behavioral recording were controlled by MATLAB (MATLAB 2017a; The MathWorks, Inc., Natick, Massachusetts) on an IBM compatible PC running on Windows 10. After completion of the discounting task, participants were thanked and fully debriefed.

**Figure 1.**
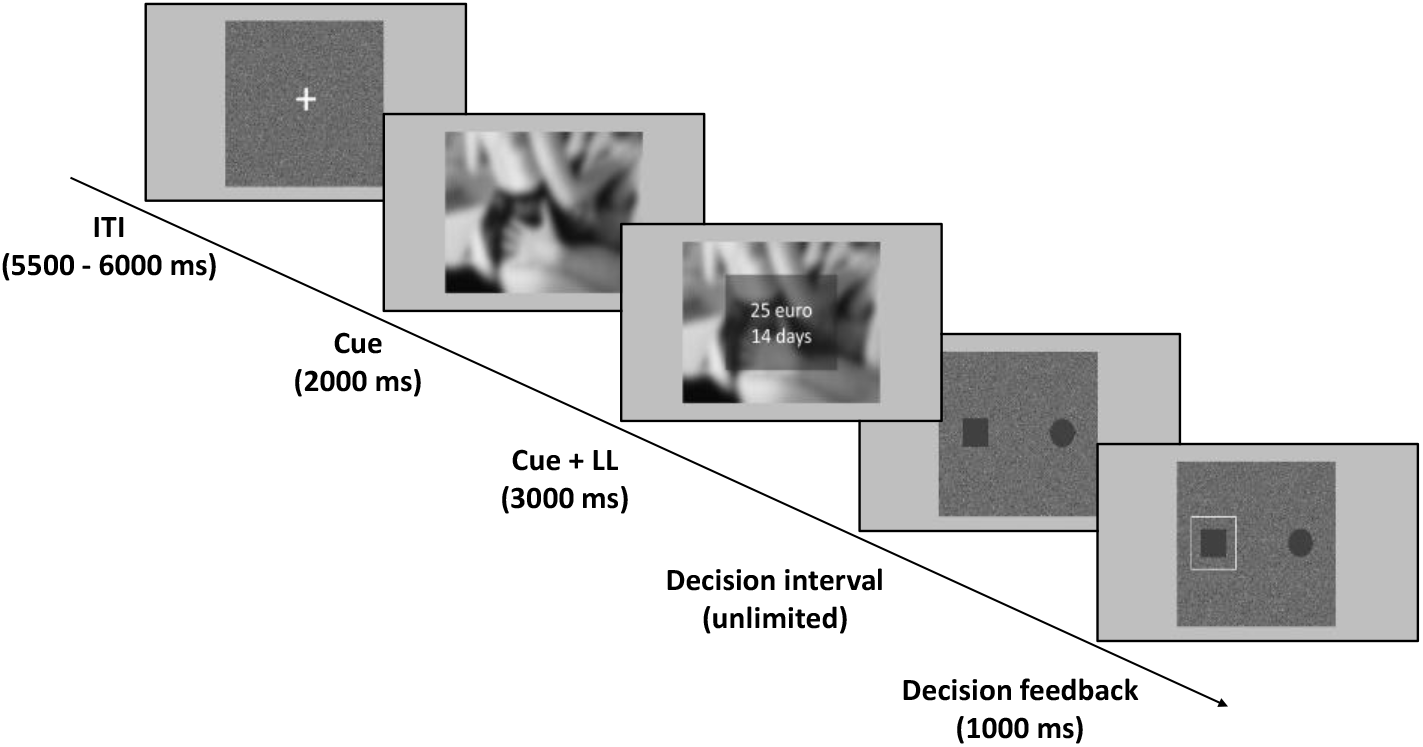
Example trial from the experimental temporal discounting task. LL = larger-later reward; ITI = intertrial-interval (onset to onset).

### 2.6. Data analysis

Data analysis was conducted using MATLAB and R (Version 3.5.1). For frequentist statistical approaches including within-subjects repeated-measures variables (e.g. rmANOVA) we report Greenhouse-Geisser-corrected *p*-values, degrees of freedom and epsilon values if the assumption of sphericity was violated. Effect sizes of significant results (*p* < 0.05) are reported as proportion of explained variance (partial eta squared). For follow-up tests, Bonferroni correction for multiple comparisons was applied.

#### 2.6.1. Analysis of choice data

To quantify temporal discounting, we used two complementary approaches. As a model-free approach we computed the area under the empirical discounting curve (AUC; Myerson et al., 2001). Our model-based approach then utilized adapted versions of the hyperbolic model (Mazur & Coe, 1987) and the softmax choice rule (Eq. 1 & Eq. 2, see above).

##### Model-free approach

Model-free approaches avoid potential issues with parameter estimation or the commitment to a specific mathematical framework (e.g. hyperbolic vs. exponential discounting). We computed the area under the empirical discounting curve (AUC) as a model free measure of discounting (Myerson et al., 2001), corresponding to the area under the connected data points that describe the decrease of the subjective value (y-axis) over time (x-axis). Each delay is expressed as a proportion of the maximum delay and plotted against normalized subjective (discounted) value as a fraction of objective value. The area of the resulting trapezoids was computed as follows (Myerson et al., 2001):

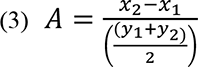

The sum across trapezoids then corresponds to the individual AUC. AUC-values were compared between neutral, aversive and erotic cue conditions using repeated measurement ANOVA. Here, smaller AUC-values indicate steeper discounting.

##### Computational Modeling

We used hierarchical Bayesian modeling to fit adapted versions of the hyperbolic model with softmax action selection. For each parameter (log*(k)* & softmax *ß*) we fit group-level distributions for the neutral condition from which individual subject parameters were drawn. To model (cue-) condition effects, we fit two separate group-level distributions modeling deviations from the neutral condition for aversive and erotic cues, respectively (“shift”-parameters, Eqs. 4 & 5) (Pedersen et al., 2017).

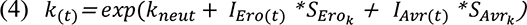

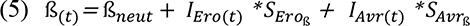

Here, *I_Ero_* and *I_Avr_* are dummy-coded indicator variables coding the respective experimental conditions (erotic vs. aversive). *S_Ero_* and *S_Avr_* are the subject-specific parameters modeling changes in log(*k*) and *ß* depending on the condition on trial *t*. These trial-wise estimates for *k* and *ß* were then used to calculate the subjective value (*SV*) of the larger-later reward (LL) as well as the probability to choose the respective option (see Eqs. 1 & 2, see above). Parameter posterior distributions were estimated via Markov Chain Monte Carlo (MCMC) as implemented in JAGS (Version 4.2) using R and the r2jags Package (Plummer, 2003). The prior distributions for the group-level parameters of the hierarchical model are listed in Table 1. JAGS model code is publicly available at OSF (https://osf.io/bk64d/). Model convergence was assessed via the Gelman-Rubinstein convergence diagnostic *R̂* and values of 1 ≤ *R̂* < 1.03 were considered acceptable. We ran 2 chains with a burn-in period of 1.280k samples and thinning factor of 2. 20k samples were then retained for further analysis. For details on MCMC convergence see Gelman and Rubin (1992).

**Table 1.**
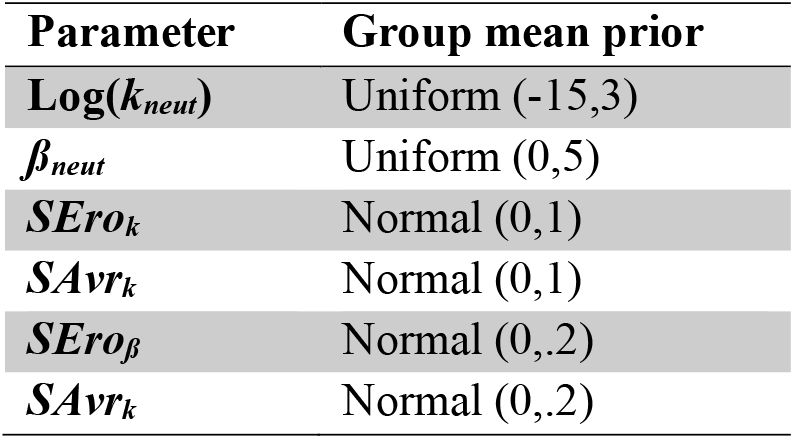
Priors of group-level parameter means. Ranges (uniform distribution), means and variances (normal distributions) were chosen to cover numerically plausible values.

We evaluated cue effects on parameter estimates using Bayesian statistics (see Kruschke, 2010). Specifically, we analysed the posterior distributions of parameters modeling group mean condition effects (as reflected in the *S_Ero_* and *S_Avr_* parameters, see Eqs. 4 & 5) by computing their highest density intervals (HDI) and assessed their overlap with zero. Furthermore, we evaluated the degree of evidence for directional effects by computing directional Bayes Factors (dBF) for erotic (*S_Ero_*) and aversive (*S_Avr_*) shift-parameters for log(*k*) and *ß*. A dBF corresponds to the ratio of the posterior mass of the shift-parameter distributions above zero to the posterior mass below zero (Marsman & Wagenmakers, 2017). Directional Bayes Factors above 3 can be interpreted as moderate evidence in favor of a positive or increasing effect while Bayes Factors above 12 are interpreted as strong evidence (Beard et al., 2016). Bayes Factors below 0.33 are likewise interpreted as moderate evidence in favor of a negative or decreasing effect on the respective parameter. As individual subject parameters were drawn from group-level distributions, and parameter estimates cannot be considered independent, we did not include a classical frequentist statistics approach (as preregistered) to assess differential cue effects on choice.

#### 2.6.2. Analysis of physiological data

##### Pupil data

Pupil data was first divided into segments ranging from 1000ms before, until 5000ms after image onset. Next, segments were screened for outliers and implausible values. Here, we slightly adjusted our preregistered pre-processing protocol to better control for putative artifacts. We defined outliers as values which exceed the respective trial mean by more than two standard deviations (Finke et al., 2017). Such outliers (2.9% of all data within relevant epochs) and missing data points due to blinks or other artefacts (4.2% of all data within relevant epochs) within the trial were linearly interpolated. Trials including more than 12% missing datapoints or outliers were excluded from further analyses (8.2% of trials on average). Subjects exhibiting more than 90% invalid trials were completely discarded from further analyses (n=2). Next, pupil data was down-sampled to 20 Hz by means of a moving average filter and median baseline-corrected. As the appearance of the choice options starting at 5s post image onset might induce changes in luminance and additional pupil responses, we calculated the mean pupil diameter for the whole image presentation interval (0-5s) excluding dilation data from the decision period (see preregistration). Further, mean pupil size was calculated in ten bins of 0.5s following image onset. Pupil size measures were compared between cue conditions using repeated measurements analysis of variance (ANOVA).

##### Heart rate data

Heart rate data were visually screened and manually corrected for major artifacts. We linearly interpolated these intervals via Biopacs built-in connect-endpoints-algorithm. We used custom MATLAB routines to detect QRS-complexes within ECG-data because Biopacs peak detection algorithm (preregistered) missed complexes in multiple subjects resulting in a strong underestimation of heart rates. Nonetheless, one subject had to be discarded due to poor data quality which resulted in an unreliable R-wave detection. To investigate phasic heart rate changes in response to erotic, aversive and neutral cues in real-time, we adopted a weighted average approach (Graham, 1978; Velden & Wölk, 1987). We used the interbeat-interval length to calculate the mean heart rate change in 0.5s bins relative to an 2s baseline prior to image onset. Trials containing implausibly low (HR < 30) or high (HR > 180) heart rate frequencies were completely discarded. Interbeat-intervals and the corresponding heart rate were weighted by their respective real time fraction in the bin. The resulting weighted means across the whole image presentation interval (0-5s) as well as within the respective time bins for all three conditions were compared using repeated measurements ANOVA.

##### Skin conductance data

We manually screened electrodermal activity data and corrected for major artifacts and signal losses. These intervals were linearly interpolated via Biopacs built-in connect-endpoints-algorithm. To achieve an improved detection and exclusion of rapid transients, drifts and other more subtle artifacts, we extended and adapted our preregistered preprocessing steps in the following way: Raw data was first low-pass filtered (1 Hz) and smoothed by a moving average filter (63 samples). Next, data was down-sampled to 62.5 Hz to reduce computational load. Instead of using Biopacs built-in analysis routine to derive phasic activity from the raw signal (preregistered) we used Ledalab’s automated continuous decomposition analysis routines to decompose skin conductance (SC) data into its tonic and phasic components which in turn reflect underlying sudomotor nerve activity (Benedek & Kaernbach, 2010a; Benedek & Kaernbach, 2010b). As sweat gland activity typically lags behind sympathetic nervous system changes, we calculated the maximum value of phasic activity and the response latency of the first significant skin conductance response (SCR) within a time window of 1-6s post image onset (significant SCR amplitude threshold of 0.03*µS*). In an exploratory approach we further assessed the total number of evoked SCR’s, the sum of amplitudes of all significant SCR’s as well as the mean phasic activity within the respective time window. The different outcome measures were compared between cue-conditions using repeated measurements ANOVA.

##### Further evaluation of physiological cue responses

After evaluation of cue effects on autonomic nervous system activity (pupil dilation, heart rate, EDA) we next explored associations amongst physiological measures on the single-subject level. For this purpose, we used the single-trial mean changes in pupil diameter, heart rate and phasic electrodermal activity to compute Pearson’s correlation coefficients. Single-trial data were standardized within subject. Using one-sample *t*-tests, we determined whether the mean of the Fisher z-transformed correlation coefficients differed from zero. Furthermore, we calculated cue-reactivity indices for each measure by computing difference scores between the mean response to erotic vs. neutral images and aversive vs. neutral images, respectively. To explore potential associations between the different physiological cue-reactivity effects, these difference scores were correlated across participants via Pearson’s correlations.

#### 2.6.3. Analysis of arousal effects on temporal discounting

##### Model-free approach

We next explored the association between the estimated erotic and aversive shift-parameters from the hierarchical model (Eqs. 4 & 5) and physiological cue-reactivity indices (see above). That is, we correlated within-subject physiological difference scores (physiological responses following erotic vs. neutral trials [erotic cue-reactivity], aversive vs. neutral trials [aversive cue-reactivity]) with model-based measures of behavioral effects (*S_Ero(k,ß)_*, *S_Avr(k,ß)_*).

##### Computational modeling

In the next step we investigated physiological arousal effects on choice behavior on the single-trial level. As cue-reactivity was most robustly associated with changes in pupil diameter (see Figure 2), we initially used within-subject standardized pupil size as a proxy for the single-trial arousal level. To quantify effects of arousal on choice behavior, we included additional parameters (*EroPupil(k,ß), AvrPupil(k,ß)*) modeling changes in log(*k)* and *ß* due to the current arousal state, over and above effects of the experimental condition, as follows (see Eqs. 6 & 7):

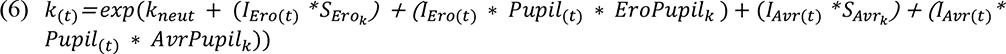

**Figure 2.**
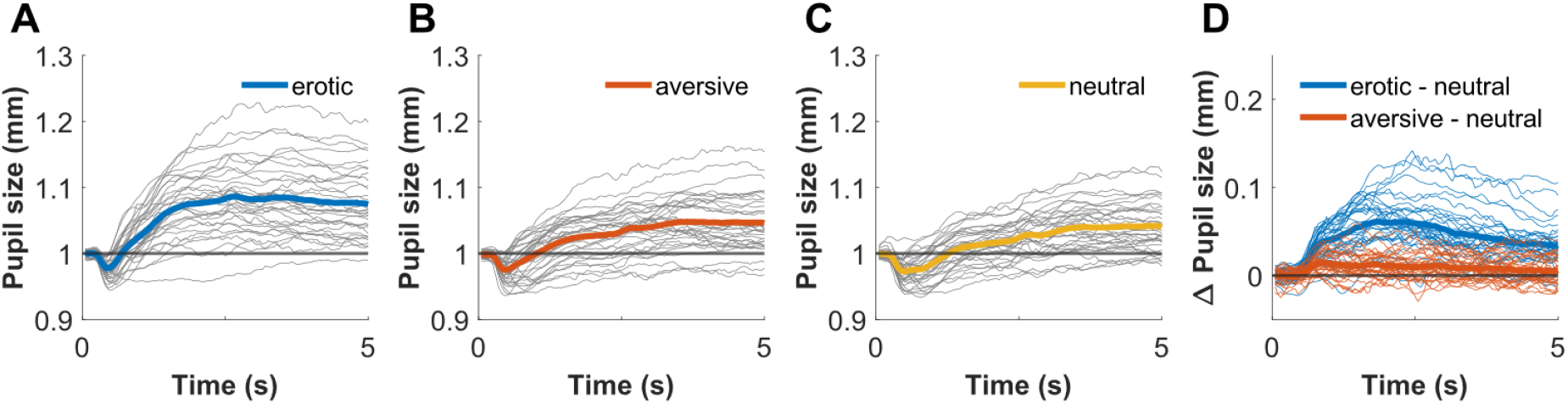
Baseline-corrected grand average pupil responses (in mm) following erotic **(A)**, aversive **(B)** and neutral **(C)** image onset; Thin gray lines depict mean single-subject pupil trajectories. Intra-individual contrasts in pupil size following image presentation are depicted in (**D**); **blue lines**: erotic – neutral; **red lines**: aversive – neutral.

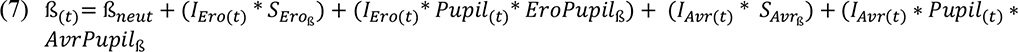

Here *I_Ero_* and *I_Avr_* again denote dummy-coded indicator variables coding the experimental condition and *S_Ero_* and *S_Avr_* are the subject-specific parameters modeling changes in log(*k*) and *ß* depending on the condition in the current trial *t*. *EroPupil* and *AvrPupil* capture additional condition-specific variation in the model parameters (log(*k*), *ß*) due to trial-wise pupil dilation. These modulated *k*- and *ß*-parameters were then used to calculate the subjective value (*SV*) of the delayed option and the respective choice probabilities. The single-subject parameters *EroPupil* and *AvrPupil* were again drawn from group-level normal distributions, with mean and variance hyper-parameters that were themselves estimated from the data. The prior distributions for the group-level parameters of the hierarchical model, the individual-level parameters as well as all relevant model equations are publicly available at OSF (https://osf.io/bdwfa/).

In a final model, we examined trial-wise arousal effects irrespective of the experimental condition, and jointly for all three physiological measures (pupil size, heart rate, EDA). Instead of fitting choice data with a logistic model on the single-subject level, using standardized mean pupil-size, heart rate and phasic electrodermal activity as regressors (as preregistered), we adapted the above-mentioned hierarchical model as follows: Erotic and aversive shift-parameters which captured condition-dependent changes in log(*k*)- and *ß* were removed from the model. Instead, we now included three arousal regressors (*PupilReg*, *EcgReg*, *EdaReg*) modeling changes in log(*k*) and *ß* as a function of trial-wise mean changes in pupil size, heart rate or skin conductance (Eqs. 8 & 9):

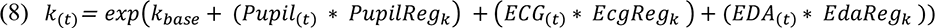

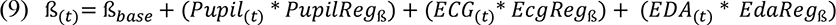

Again, priors and model code are publicly available at OSF (https://osf.io/ajxkd/).

#### 2.6.4. Data and code availability

Behavioral and physiological data as well as JAGS model code is available on the Open Science Framework (https://osf.io/dtwg3/files).

## 3. Results

### 3.1. Physiological cue responses

Analysis of the physiological data proceeded in the following steps. First, we examined cue-evoked changes in autonomic nervous system activity. We did this separately for each physiological measure (pupil size, heart rate, EDA). Next, we used Pearson’s correlations to assess the associations between evoked physiological responses following image onset on the single-trial level as well as the concordance between differential responses to erotic and aversive image content (difference scores, see above). Unless otherwise stated, all analyses were conducted as preregistered (https://osf.io/swp4m/).

#### 3.1.1. Pupil diameter change

Evoked changes in pupil size following affective image presentation are depicted in Figure 2. After an initial dip reflecting a small initial light reflex (Beatty & Lucero Wagoner, 2000) at around 0.5s post image onset, we observed a substantial pupil dilation response, differently pronounced for erotic, aversive and neutral pictures (Figure 2 (**A, B & C**)). A two-factor repeated measures ANOVA (within factors: image condition, time bin (0.5s/bin)) showed a significant main effect for image condition (*F*[2,64] = 91.17, *p* < .001, *η*_p_^2^= 0.74, *ε* = 0.08). Post-hoc t-tests indicated increased pupil dilation responses following aversive compared to neutral stimuli, (*t*_(aversive, neutral)_ = 4.49, *p* < .001, CI_Diff(95%)_ = [0.005;0.01]), a pattern which was even more pronounced for erotic stimuli (*t*_(erotic, neutral)_ = 10.47, *p* < .001, CI_Diff(95%)_ = [0.03;0.05]; *t*_(erotic, aversive)_ = 9.36, *p* < .001, CI_Diff(95%)_ = [0.03;0.04].

Furthermore, we observed a significant interaction effect of the factors image condition and time bin (*F*[18,576] = 43.30, *p <* .001, *η*_p_^2^= 0.58, *ε* = 0.08). As illustrated in Figure 2 (**D**), condition-dependent differences in pupil size emerged between the first and second bin, increased until the fifth bin and then slightly decreased over time. All pairwise comparisons are depicted in Table S1 (Supplementary Materials). Intra-individual contrasts indicated that in particular the increased pupil responses to erotic stimuli were highly consistent between subjects (Figure 2 (**D**)).

#### 3.1.2. Heart rate change

Heart rate decelerated in response to image onset irrespective of image condition, likely reflecting an initial orienting response (Hare, 1972). In accordance with previous studies, the deceleration pattern was numerically most prominent for erotic and aversive image conditions (Figure 3 (**A, B & D**); Abercrombie et al., 2008). However, a repeated measurements ANOVA (within factors: image condition, time bin (0.5s/bin)) did not reveal a significant main effect for image condition (*F*[2,66] = 1.74, *p* = .196, *ε* = 0.11). The interaction of the factors image condition and time bin indicated differences in heart rate deceleration at trend level across timepoints (*F*[18, 594] = 3.00, *p =* .052, *ε* = 0.11), which were most pronounced 3-3.5s post image onset.

**Figure 3.**
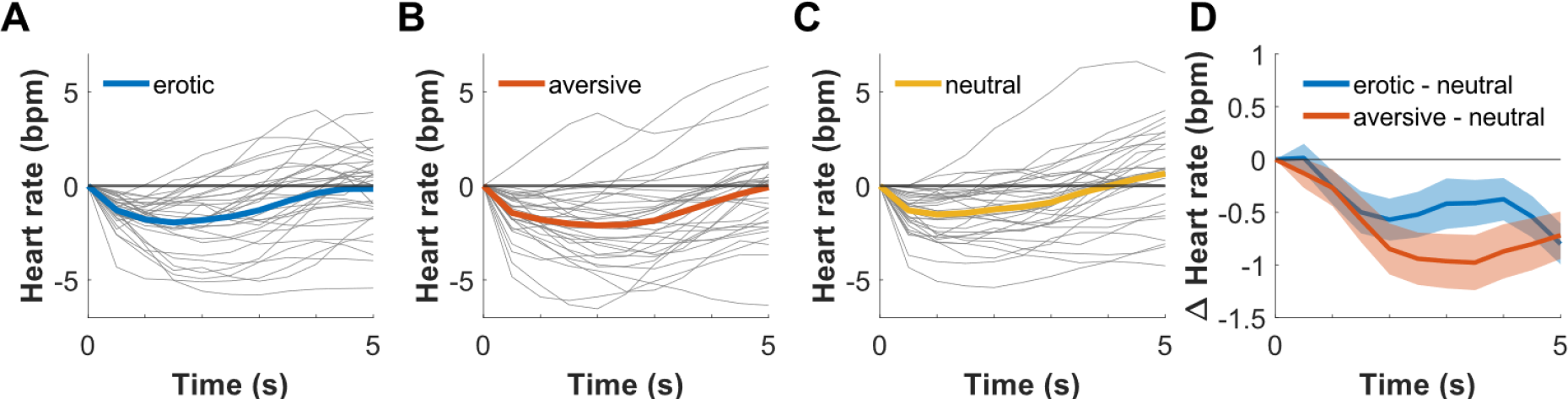
Baseline-corrected grand average heart rate changes (in bpm) following erotic (**A**), aversive (**B**) and neutral (**C**) image onset; Thin gray lines depict single-subject heart rate trajectories; Grand average contrasts of average heart rate change are shown in (**D**); **blue line**: erotic – neutral; **red line**: aversive – neutral. Shaded areas depict standard errors (SE).

#### 3.1.3. Electrodermal activity change

To evaluate possible cue effects on alterations in electrodermal activity we assessed the latency of the first evoked skin conductance response (SCR) as well as the maximum phasic peak following image presentation (Figure 4 (**A, B**)). Contrary to our expectations we did not find differential effects of image condition on those measures (Latency (SCR): *F*[2,68] = 0.13, *p =* .876; Phasic activity: *F*[2,68] = 1.41, *p* = .252).

**Figure 4.**
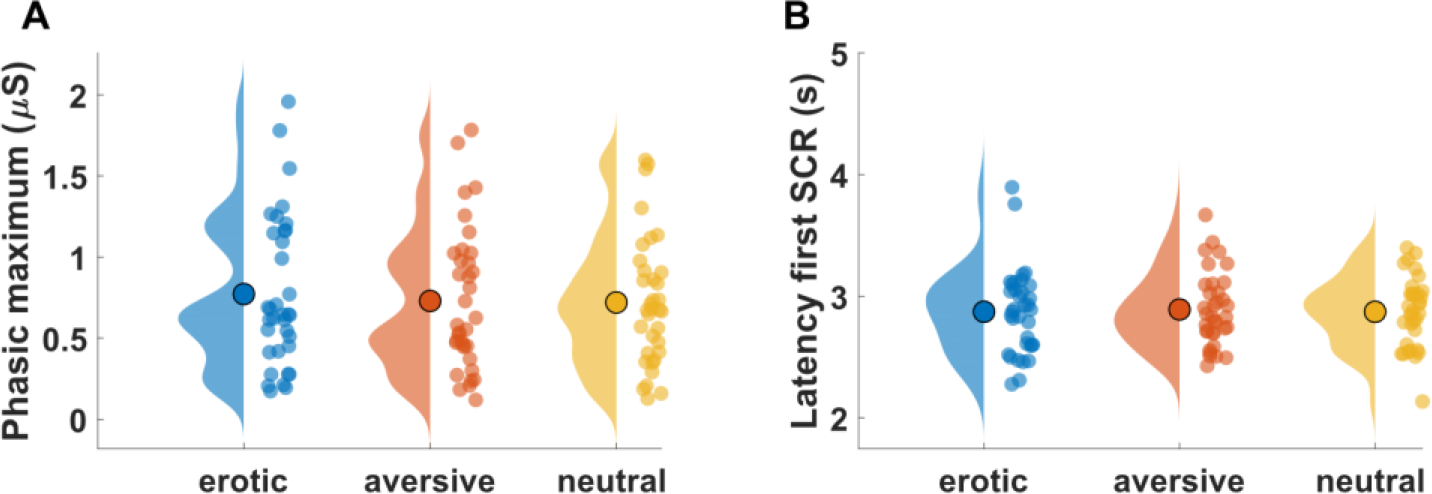
Maximum value of phasic activity **(A)** and latency of first skin conductance response (SCR) **(B)** in the interval 1-6 seconds post image onset. Colored dots depict single-subject means. *µS* = microsiemens.

In an exploratory analysis (not preregistered) we extracted three additional measures to quantify cue-evoked electrodermal responses and compared them between image conditions: the sum of SCR amplitudes, the number of SCR’s, as well as mean phasic activity within the interval 1-6s post image onset. Sum of SCR amplitudes and mean phasic activity did not differ between conditions, whereas repeated measures ANOVA indicated significant differences in the number of SCR’s between conditions (F[2,68] = 4.17, *p* = .020, *η*_p_^2^= 0.11). There was a greater mean number of SCR’s following erotic images (mean ± Std = 83.31 ± 33.13) compared to aversive (mean ± Std = 77.83 ± 28.96) and neutral (mean ± Std = 78.06 ± 31.10). However, none of the pairwise comparisons survived Bonferroni correction. Cue effects on all three exploratory measures are shown in Figure S3 (Supplementary Materials).

#### 3.1.4. Associations amongst physiological measures

We next conducted an exploratory analysis to examine associations amongst the three physiological measures using two complementary approaches. First, we used the single-trial mean changes in pupil diameter, heart rate and phasic electrodermal activity to compute Pearson’s correlation coefficients. Then, concordance between differential responses to erotic and aversive image content was assessed between all three physiological measures. These analyses revealed if anything small associations (range (*r*): -.03 -.26). A detailed description of all conducted correlational analyses and respective results are depicted in Figure S4 and Figure S5 (Supplementary Materials).

### 3.2. Cue effects on temporal discounting

Having thus confirmed trial-wise changes in physiological arousal, we next quantified condition-specific changes in temporal discounting. We used model-free and computational modeling approaches, as preregistered (https://osf.io/swp4m/).

#### 3.2.1. Model-free approach

Applying repeated measures ANOVA on the area under the empirical discounting curve (AUC) revealed no significant differences between erotic (mean (AUC) = .69), aversive (mean (AUC) = .68) and neutral (mean (AUC) = .68) conditions (F[2,68] = 0.29, *p* = .753; Figure 5).

**Figure 5.**
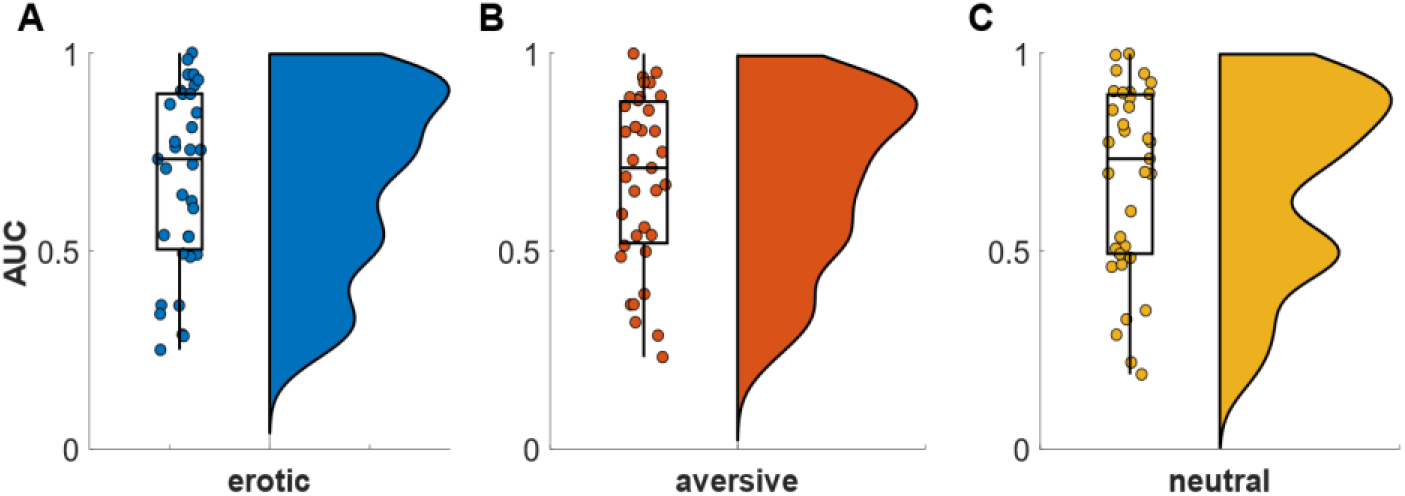
Area under the empirical discounting curve (AUC) for erotic (**A**), aversive (**B**) and neutral (**C**) cue conditions. Dots depict single-subject AUCs (means).

#### 3.2.2. Computational modeling

Using hierarchical Bayesian modeling, we fit adapted versions of the hyperbolic model with softmax action selection to the choice data. To estimate changes in discounting behavior due to erotic or aversive cue exposure, we fit group-level distributions for the neutral condition from which individual subject parameters were drawn. Subject-specific *S_Ero_* and *S_Avr_* parameter (Eqs. 4 & 5) then modeled trial-wise condition-specific changes in log(*k*) and *ß*. Examination of the posterior distributions of *S_Ero(k)_* and *S_Avr(k)_* from the computational model suggested a small but consistent decrease in discounting following erotic stimuli (95.01% of posterior distribution of *S_Ero(k)_* fell below zero). The associated directional Bayes Factor (dBF) yielded dBF = 19.78, indicating that a reduction in discounting following erotic cues was 19.78 times more likely than an increase, given the data. In contrast, *S_Avr(k)_* showed a substantial overlap with zero (dBF = 0.27) (Figure 6 (**A, B**)). Further, *ß*-parameters slightly decreased following aversive cues, reflecting elevated decision noise. Inspection of the posterior distribution showed that 97.16% of the *S_Avr(ß)_* posterior distribution fell below zero. The directional Bayes Factor (dBF = 40.37) indicated that an increase in decision noise following aversive stimuli was 40.37 times more likely than a decrease, given the data. Such an effect was not observed for erotic cues (dBF = 0.71) (Figure 6 (**D, E**)).

**Figure 6.**
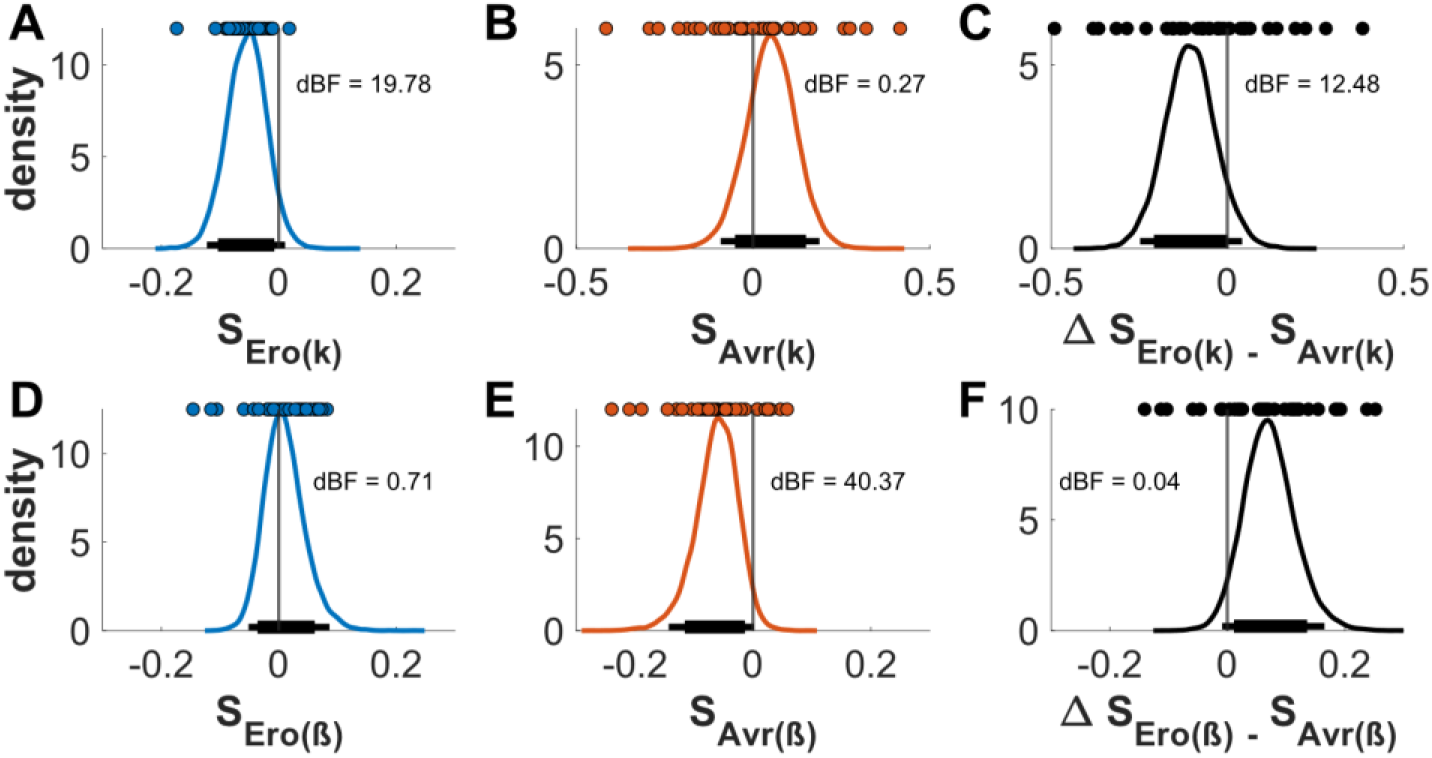
Posterior distributions for erotic (*S_Ero_*; blue) and aversive (*S_Avr_*; red) shift-parameters as well as their differences (black); **A-C**: Log(*k*); **D-F**: Softmax (*ß*); Colored dots depict single-subject means. Thick and thin horizontal lines indicate 85% and 95% highest density intervals; dBF = directional Bayes Factor.

To validate model parameters, we explored associations between *S_Ero(k)_* and *S_Avr(k)_* and model-free measures (AUC-values (3.2.1), larger-later choice proportions). Correlations between model parameters and model-free measures were consistently in the expected direction (see Figure S6, Supplementary Materials).

### 3.3. Arousal effects on temporal discounting

Both parameters from the computational model (*S_Ero_*, *S_Avr_*) and pupil responses were affected by cue condition (although in the case of log(*k*) these effects were in the opposite direction as predicted). We next explored whether these effects were associated on the subject-level. We extracted the means of the posterior distributions of *S_Ero_*-, and *S_Avr_*-parameters per participant and computed Pearson’s correlations between those means and the difference scores between the average pupil response to affective (erotic & aversive) and neutral stimulus material (exploratory analysis). These analyses revealed overall small and non-significant associations (see Figure S7, Supplementary Materials).

We next examined whether the single-trial arousal predicted trial-wise changes in temporal discounting (exploratory analysis). As in particular pupil responses reliably differentiated between cue conditions (see Figure 2), we initially focused on this measure. Arousal level was quantified via mean single-trial pupil dilation and evaluated in terms of its modulating effect on discounting behavior. Specifically, we set up an additional hierarchical Bayesian model in which trial-wise parameters (log(*k*) and *ß*) were allowed to vary both according to the cue condition (*S_Ero(k,ß)_, S_Avr(k,ß)_*) and according to the trial-wise arousal level as reflected in pupil dilation responses (*EroPupil_(k,ß)_, AvrPupil_(k,ß)_*; see Eqs. 6 & 7; exploratory analysis). We reproduced the attenuation of log(*k*) in the erotic condition (see Figure 6 (**A**)) also in this model. As can be seen from Figure 7 (**A**), 91.34% of the posterior distribution for *S_Ero(k)_* fell below zero. The associated dBF of 11.62 indicated some evidence for a decreasing effect of the erotic cue condition on log(*k*) which was independent of the trial-wise arousal level. However, this effect was somewhat less pronounced than in the model without a pupil predictor (Figure 6). This was due to the fact that trial-wise pupil dilation (*EroPupil_(k)_*) also exhibited a small negative effect on log(*k*) (Figure 7 (**B**), dBF = 8.70, 88.29% of the posterior distribution fell below zero). Although this effect was numerically small, it was consistent across subjects (Figure 7 (**B**)). The posterior distribution for *S_Avr(k)_* showed substantial overlap with zero (Figure 7 (**C**), dBF = 0.57). Likewise, the effect of pupil dilation following aversive cues (*AvrPupil_(k)_*) was of inconclusive directionality (dBF = 0.43).

**Figure 7.**
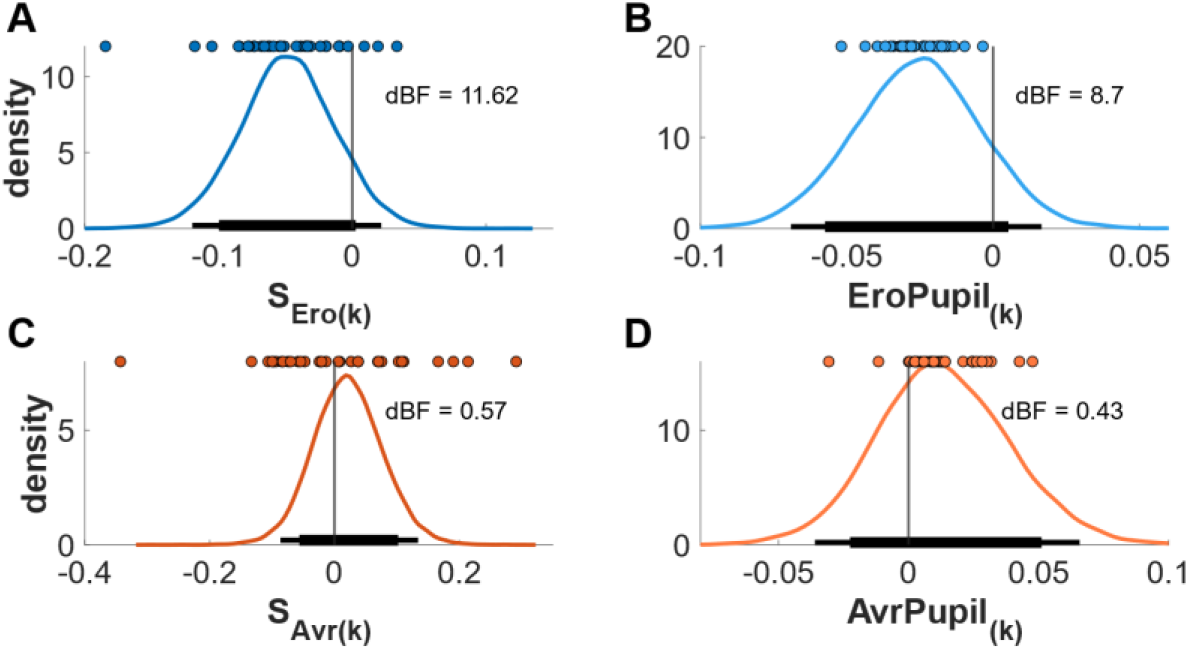
Posterior distributions for erotic (*S_Ero(k)_)* and aversive (*S_Avr(k)_)* shift parameters on log(*k)* (**A, C**) and shift parameters due to single trial arousal state following erotic (*EroPupil_(k)_*) and aversive (*AvrPupil_(k)_*) stimuli (**B, D**). Colored dots depict single subject means. Thick and thin horizontal lines indicate 85% and 95% highest density intervals; dBF = directional Bayes Factor.

Posterior distributions of the *S_Ero(ß)_*-parameter (Figure 8 (**A**)) indicated that, if anything, decision noise decreased (softmax (*ß*) increased) in response to erotic stimuli while it increased in response to aversive image content (*S_Avr(ß);_* Figure 8 (C), dBF*_(SEro(ß))_* = 0.05; dBF_(*SAvr(ß))*_ = 5.88). The effect of single-trial arousal state on decision noise in the erotic condition (*EroPupil_(ß)_*) was of inconclusive directionality (Figure 8 (**B**), dBF = 2.09), whereas, if anything, this association was positive for aversive cues (*AvrPupil_(ß)_*; Figure 8 (**D**), dBF = 0.28).

**Figure 8.**
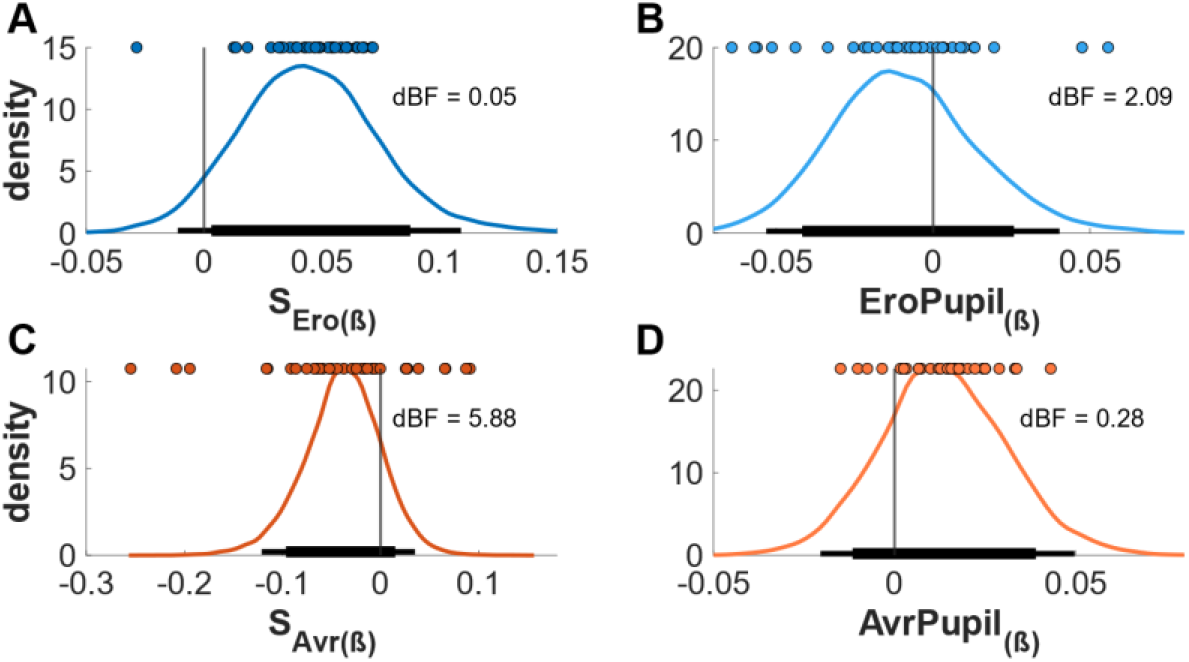
Posterior distributions for erotic (*S_Ero(ß)_)* and aversive (*S_Avr(ß)_)* shift-parameters on *ß* (**A, C**) and shift-parameters due to single trial arousal state following erotic (*EroPupil_(ß)_*) and aversive (*AvrPupil_(ß)_*) stimuli (**B, D**). Colored dots depict single subject means; Thick and thin horizontal lines indicate 85% and 95% highest density intervals; dBF = directional Bayes Factor.

We next assessed arousal effects on temporal discounting irrespective of overall condition effects (preregistered analysis). Single-trial mean pupil diameter, heart rate and phasic electrodermal activity were included in the trial-wise computation of log(*k*) and *ß* (see Eqs. 8 & 9), yielding separate estimates of the effects of each physiological measure on log(*k*) and *ß*. Posterior distributions for all three effects are depicted in Figure 9. Whereas trial-wise pupil response and heart rate change were both, if anything, associated with decreases in discounting (dBF*_PupilReg(k)_* = 17.98, dBF*_EcgReg(k)_* = 3.41), phasic electrodermal activity showed no systematic association with log(*k*) (dBF*_EdaReg(k)_* = 0.92; see Figure 9 (**A-C**)). In contrast, whereas single-trial heart rate change was negatively associated with decision noise (dBF*_EcgReg(ß)_* = 9.27), pupil diameter and phasic electrodermal activity were not systematically associated with *ß* (Figure 9 (**D-F)**; dBF*_PupilReg(ß)_* = 0.37; dBF*_EdaReg(ß)_* = 0.46).

**Figure 9.**
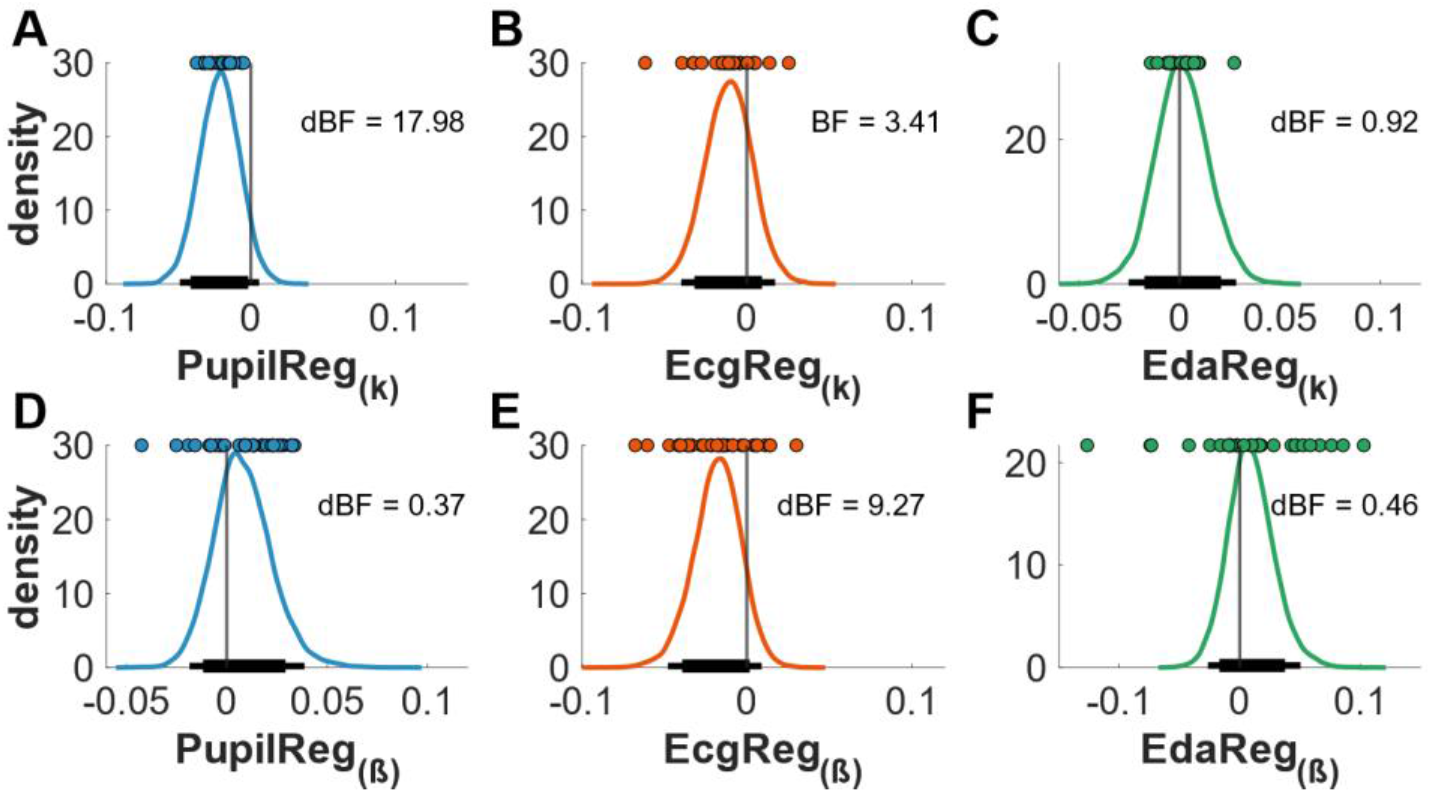
Posterior distributions depicting single trial pupil (**A & D**) heart rate (**B & E**) and skin conductance (**C & F**) effects on choice. Colored dots depict single subject means. Thick and thin horizontal lines indicate 85% and 95% highest density intervals; dBF = directional Bayes Factor.

As effects of physiological measures on log(*k*) and *ß* might be affected by habituation, we investigated cue-evoked physiological responses over the course of the experiment. To this end, trials of pupil diameter, heart rate and phasic electrodermal activity data, were binned into three time-bins separately for neutral, erotic and aversive cue conditions. There was overall limited evidence for habituation across measures (Figure S8, Supplementary Materials). For example, cue-evoked pupil dilation significantly decreased over time, but this effect was numerically small, and pupil dilation still differentiated reliably between cue conditions across all phases of the experiment. Heart rate and electrodermal activity data also showed small but non-significant reductions in evoked responses.

### 3.4. Physiological and behavioral indices of cognitive effort

We finally hypothesized that difficult trials (high decision conflict, i.e. high cognitive effort) would result in a more pronounced pupil dilation across conditions. This effect should be most pronounced following LL-onset and might last until a decision is made. Here, we slightly deviated from our preregistered analysis plan (analysis of trial difficulty- and cue-condition effects on pupil dilation in two separate general linear models (GLMs; https://osf.io/swp4m/)). Instead, we used three complementary approaches. First, we compared pupil size starting at LL-onset until decision screen onset between easy, medium and hard trials using repeated measures ANOVA. Second, we used a linear mixed model to analyze whether median pupil size in the interval one second prior to the response differed between trial difficulty levels. Third, using the Pupil Response Estimation Toolbox (PRET; Denison et al., 2020) we constructed a general linear model (GLM) to examine trial difficulty- and cue-effects on pupil dilation.

Contrary to our expectations, pupil size was not significantly modulated by decision conflict (see Supplementary Materials for details on conducted *<u>analyses</u>* and *<u>results</u>*). In an exploratory approach, we also investigated whether response times (RTs), which are typically longer under high decision conflict, changed as a function of trial difficulty (Peters & Büchel, 2010) or cue condition. Results from a linear mixed model indeed indicated longer RTs for medium and hard trials compared to easy trials, while RTs did not differ significantly between hard and medium trials. There were no further significant main effects or interactions. We also explored whether behavioral (RTs) and physiological (pupil dilation) indices of cognitive effort showed associations at the subject level. There were no significant associations (see Supplementary Materials for details on conducted *<u>analyses</u>* and *<u>results</u>*).

## 4. Discussion

Here we investigated the effects of erotic, aversive and neutral visual cues on temporal discounting (TD) in a trial-wise design. We used comprehensive monitoring of autonomic nervous system (ANS)-activity in response to cue presentation to assess contributions of trial-wise physiological arousal to TD modulations. Physiological arousal was robustly elevated following aversive and in particular erotic cue exposure. Contrary to our predictions, if anything, we observed evidence for a small *decrease* in TD following erotic cues. In contrast, aversive cues tended to increase decision noise. Trial-wise arousal only accounted for minor variance over and above aversive and erotic condition effects.

### 4.1. Cue effects on autonomic activity

Autonomic activity was assessed using three complementary measures (pupil size, heart rate, electrodermal activity) which are differentially affected by sympathetic and parasympathetic branches of the nervous system (Bradley et al., 2008). Whereas changes in skin conductance responses are mainly driven by sympathetic activity (Dawson et al., 2007; Posada-Quintero et al., 2016; Venables & Christie, 1980), modulations of heart rate (Berntson et al., 1997) as well as pupil size (Fotiou et al., 2000; Löwenfeld, 1999; Steinhauer et al., 2004) result from an interplay of parasympathetic and sympathetic afferents.

Cue-evoked arousal modulations were successfully captured by pupil dilation. A small onset-related pupil constriction was followed by a long-lasting dilation response which was substantially increased for both erotic and aversive cues, confirming previous findings of appetitive (Finke et al., 2017) and aversive stimulus processing (Kinner et al., 2017). A direct comparison confirmed previous findings of increased responses to erotica (Bradley et al., 2008; Bradley & Lang, 2015; Henderson et al., 2014). Such dilatory pupil responses have been observed following various cues with high motivational and behavioral relevance, including engaging sounds (Partala et al., 2000), task-relevant stimuli (Kahneman & Beatty, 1966) and surprising events (Preuschoff et al., 2011), which have been closely linked to phasic activations of locus coeruleus (LC) neurons (Aston-Jones & Bloom, 1981b, but see Aston-Jones & Cohen, 2005) and concomitant norepinephrine (NE) release (Abercrombie et al., 1988). Increased pupil dilation following both appetitive and aversive cues in our study might therefore reflect phasic modulations of the LC-NE-system.

Pupil dilation can be traced back to sympathetic and parasympathetic inputs (Schuman et al., 2020). Whereas pupil constriction is controlled via parasympathetic innervation of the pupillae sphincter, pupil dilation is controlled by sympathetic afferents to the dilator pupillae muscle of the iris (Andreassi, 2000; Loewenfeld, 1999). In our data, we observed no substantial cue-effects on the initial pupil constriction. However, late dilatory responses clearly discriminated between conditions, suggesting that this effect might be mediated by sympathetic involvement.

Heart rate (HR) was also modulated by cue type. A short-latency HR-deceleration was most pronounced for aversive and erotic cues. Such vagally mediated HR-suppression is assumed to reflect an initial orienting response (OR; Hare, 1972), denoting an epoch of increased sensory receptivity and deepened encoding during pleasant and unpleasant stimulus perception (Abercrombie, 2008). In line with previous literature (Bradley et al., 2001; Jönsson et al., 2008), our data indicate that such deceleration patterns appeared to be more stable following aversive stimuli. However, although the observed HR-responses might reflect modulations of parasympathetic nervous system activity by erotic and aversive cues, moderately high single-subject variance prevented significant condition differences.

Contrary to our pre-registered hypotheses, participants exhibited no overall increase in skin conductance response (SCR) amplitudes following erotic or aversive images compared to neutral. Exploratory analyses revealed largest SCR-amplitudes following erotica in the first third of the experiment, an effect that substantially habituated over time, thereby reducing overall condition differences (see Supplementary Materials). SCR’s are known to be sensitive to habituation (Steiner & Barry, 2014). In addition to SCR-amplitudes, the number of evoked SCR’s following erotic images was numerically (but not significantly) increased compared to aversive and neutral conditions. However, contrasting with prior studies (Kinner et al., 2017; Vujovic et al., 2014), participants showed no increased electrodermal responsiveness to aversive image content. Although SCR’s are a well-established measure of ANS-activity in response to pleasant and aversive stimuli (Bernat et al., 2006; Christopoulos et al., 2019), sex differences in affective picture processing have been observed (Bradley et al., 2001). Women exhibit greater physiological reactivity to aversive material compared to men (Chentsova-Dutton & Tsai, 2007; Lithari et al., 2010). In contrast, SCR’s in men are largest in response to erotic cues (Bradley et al., 2001). The fact that we only recruited male participants could therefore account for the lack of SCR modulation by aversive stimuli.

In sum, we found considerable evidence that arousal (pupil size, heart rate) was successfully modulated by our experimental conditions.

### 4.2. Cue effects on temporal discounting

Although exposure to erotic and aversive stimuli induced substantial changes of physiological arousal (pupil size, heart rate), we did not observe strong modulations of temporal discounting (TD) following cue exposure. Whereas model-free measures of impulsivity (AUC, see Methods section) did not indicate significant condition effects, posterior distributions of condition effects on log(*k*) from the computational model (see Eq. 4) suggested a small but consistent *decrease* in discounting following erotica (dBF = 19.78). This discrepancy between model-free and model-based measures might be due to the fact that AUCs were estimated separately for erotic, aversive and neutral conditions while single-subject parameter changes in the computational approach were modeled via additive shift-parameters that allowed deviations from a baseline condition (neutral) for aversive and erotic cues, respectively, potentially increasing sensitivity to subtle within-subject effects.

Earlier studies found that exogeneous cues might modulate TD (Herman et al., 2018). More specifically, block-wise presentation of appetitive (Li, 2008) and especially erotic cues (Kim & Zauberman, 2013; Van den Berg et al., 2008; Wilson & Daly, 2004) prior to TD tasks increased discounting. This has been attributed to an out-of-domain wanting for immediate pleasure (present orientation) in response to primary reinforces like erotic picture stimuli (Van den Bergh et al., 2008). Such effects might be mediated by an elevated dopaminergic tone in reward-related brain areas following sustained presentation of highly appetitive rewards like erotic stimuli (O’Sullivan et al., 2011; Redouté et al., 2000). This matches previous evidence showing that pharmacological manipulation of dopaminergic neurotransmission can directly influence discounting behavior (Petzold et al., 2019; Pine et al., 2010; Wagner et al., 2020) though overall, the corresponding human literature is small and heterogeneous (D’Amour-Horvat & Leyton, 2014).

In stark contrast, studies investigating trial-wise cue effects on TD report highly mixed results. Studies observed increased (Guan et al., 2015; Sohn et al., 2015), or decreased (Luo et al., 2014) discounting following negative primes, and increased (Sohn et al., 2015) or unaltered (Simmank et al., 2015) discounting in response to erotic cues.

Reasons for this high variability could be manifold. Multiple mechanisms might contribute to trial-wise cue effects, potentially with opposite directionality. Animal studies suggest that both highly arousing appetitive and aversive stimuli induce a graded release of noradrenaline in cortex (NE; Ventura et al., 2008). In humans, highly arousing cues of either valence increase pupil dilation (Finke et al., 2017; Kinner et al., 2017), a measure associated with locus coeruleus (LC) activity (Aston-Jones & Cohen, 2005). NE agonists might reduce several forms of impulsivity (Robinson et al., 2008) and directly increase the preference for larger-later rewards (Bizot et al., 2011). Further, Yohimbine, an α_2_-adrenergic receptor antagonist that increases NE release, reduced discounting in humans (Hermann et al., 2019; Schippers et al., 2016). Short-term increases in NE release via erotic or aversive stimuli might therefore foster more patient choice patterns.

At the same time, stimulus-evoked physiological arousal is inextricably linked to emotional processing (Hermann et al., 2018). Various studies observed increased TD following negative emotional priming (Guan et al., 2015; Lerner et al., 2013; Moore et al., 1976) – findings that correspond to the idea that experiencing emotional distress might foster desire for immediate pleasure and reward (Tice et al., 2001). Other studies report that positive stimuli, imagining positive future events or a positive mood state can reduce discounting (Guan et al., 2015; Liu et al., 2013; Rösch et al., 2021; Weafer et al., 2013). Although these findings do not remain unchallenged (Luo et al., 2014; Simmank et al., 2015), they might support the notion of opposing valence-driven cue effects on TD. Such processes might also have contributed to the small and inconsistent findings regarding erotic and aversive cue effects on decision-making in the present study. Whereas single-trial changes in noradrenergic activity might have reduced the urge for immediate reward following both, erotic and aversive stimuli, emotional processing in response to aversive image onset might have selectively increased wanting of immediately available rewards.

Further, when comparing results across trial-wise cue-exposure studies, it is crucial to consider event sequences within single trials, which might affect value integration and choice preferences. Sohn and colleagues (2015) investigated emotional arousal effects on TD using positive, negative and neutral cues. On every trial, two pictures of the same category were presented, followed by the presentation of the smaller-sooner (SS) and larger-later (LL) rewards. Similarly, Guan and colleagues (2015) first displayed negative (arousing), neutral or happy primes. The SS- and LL-options appeared both shortly after. In contrast, in the present study, cues were presented alone for two seconds, and subsequently the LL-option was superimposed. The immediately available SS-option (20 €) was never shown throughout the experiment (Kable & Glimcher, 2007; Peters & Büchel, 2009). Therefore, neural coding of the LL-reward (Strait et al., 2015; Tricomi & Lempert, 2015) may have been selectively increased by preceding erotic cues, and decreased by aversive cues. Further work is required to examine the impact of such experimental design choices on cue-exposure effects.

We observed robust cue-evoked changes in arousal, in particular for pupil dilation (see above). However, modeling revealed that trial-wise arousal had if anything small effects on log(*k*) and *ß*. Small attenuating effects of erotic cues on log(*k*) persisted, even when trial-wise pupil dilation was included in the model. Although the erotic effect was attenuated, this was not the case for the aversive cue effects, arguing against a general valence-independent effect of physiological arousal on discounting. In a final control model, we tested whether single-trial physiological measures affected log(*k*), irrespective of overall condition effects. Confirming previous results, posterior distributions indicated if anything small (decreasing) effects of pupil- and heart rate-regressors on the discount rate.

Aversive cues increased decision noise (decreased *ß*), reflecting a reduced impact of value differences on choices. This could be attributable to distraction (Dolcos & McCarthy, 2006; Stout et al., 2018) and/or increased demands for emotion regulation (Dolcos & Mc Carthy, 2006; Ochsner et al., 2004), impeding successful value integration. However, unaltered decision noise following highly arousing erotic stimuli, again argues against an underlying arousal-driven effect.

The present study has several limitations that need to be acknowledged. First, although our data indicate alterations of ANS-activity following erotic and aversive cues (Wang et al., 2018), we did not directly assess subjective arousal. However, psychophysiological ANS-measures and subjective measures tend to co-vary (Aguado et al., 2018; Lang et al., 1990). Moreover, pupil dilation during mental imagery covaries with subjective arousal (Henderson et al., 2018). We also conducted a pilot study where image characteristics of the stimulus set were pre-rated by an independent sample. The obtained ratings suggest that erotic and aversive stimuli modulated subjective arousal. However, future studies might complement physiological recordings in response to affective cue presentation by self-reported arousal. Second, we focused on male participants because erotic cue effects on TD have been primarily examined in male subjects (Kim & Zauberman, 2013; Van den Bergh et al., 2008; Wilson & Daly, 2004). Men and women might differ in their neurophysiological reactivity to affective stimulus material and emotional processing (Bradley et al., 2001; Lithari et al., 2010; Wrase et al., 2003), and future studies should extend the present approach and include participants from both sexes.

Taken together, while appetitive and aversive cues caused substantial modulation of the physiological arousal state, behavioral effects on temporal discounting were small. If anything, highly appetitive erotic cues reduced discounting, whereas aversive cues increased decision noise. Previous trial-wise cue-exposure effects on discounting were mixed, but physiological arousal was not explicitly controlled. Using extensive computational modeling and physiological monitoring, we found no strong evidence for a major influence of trial-wise physiological arousal levels on temporal discounting.

## Acknowledgements

We are grateful to Erik Lang for helping with recruitment and data collection. This work was supported by Deutsche Forschungsgemeinschaft (PE1627/5-1 to J.P.). There are no conflicts of interest.

## Author contributions

Conceptualization: J.P., K.K.; Data curation: K.K.; Formal analysis: K.K.; Funding acquisition: J.P. Investigation: K.K.; Methodology: J.P., K.K.; Project administration: K.K.; Writing – original draft: K.K.; Writing – review & editing: J.P., K.K.

## Supplementary Materials

### Pilot study ratings of stimulus material used in the study

We used repeated measures analysis of variance (rmANOVA) to compare image ratings (arousal, valence) between conditions. Results showed that arousal ratings clearly differed between conditions (*F*[2,190] = 9961.70; *p <* .001; *η*_p_^2^ = 0.99; ε = 0.61; Figure S1). Post-hoc ttests revealed, that arousal levels were comparable for erotic and aversive cues (*t*_(erotic, aversive)_ = −1.59; *p* = .346; CI_Diff(95%)_ = [-0.14;0.02]) but differed for neutral image condition (*t*_(erotic, neutral)_ = 188.04; *p* < .001; CI_Diff(95%)_ = [5.33;5.45]); *t*_(neutral, aversive)_ = −92.49; *p* < .001; CI_Diff(95%)_ = [-5.57;-5.34]).

Valence ratings also differed between cue conditions (*F*[2,190] = 1720.40; *p <* .001; *η*^2^ = 0.95; ε = 0.87; Figure S1). Post-hoc ttests showed, that erotic and aversive images differed in their rated valence (*t*_(erotic, aversive)_ = 51.13; *p* < .001; CI_Diff(95%)_ = [4.88;5.27]). Neutral images differed from both erotic and aversive image conditions (*t*_(erotic, neutral)_ = 27.76; *p* < .001; CI_Diff(95%)_ [1.80;2.08]; *t*_(neutral, aversive)_ = 34.76; *p* < .001; CI_Diff(95%)_ = [2.96;3.32]).

**Supplementary Figure S1.**
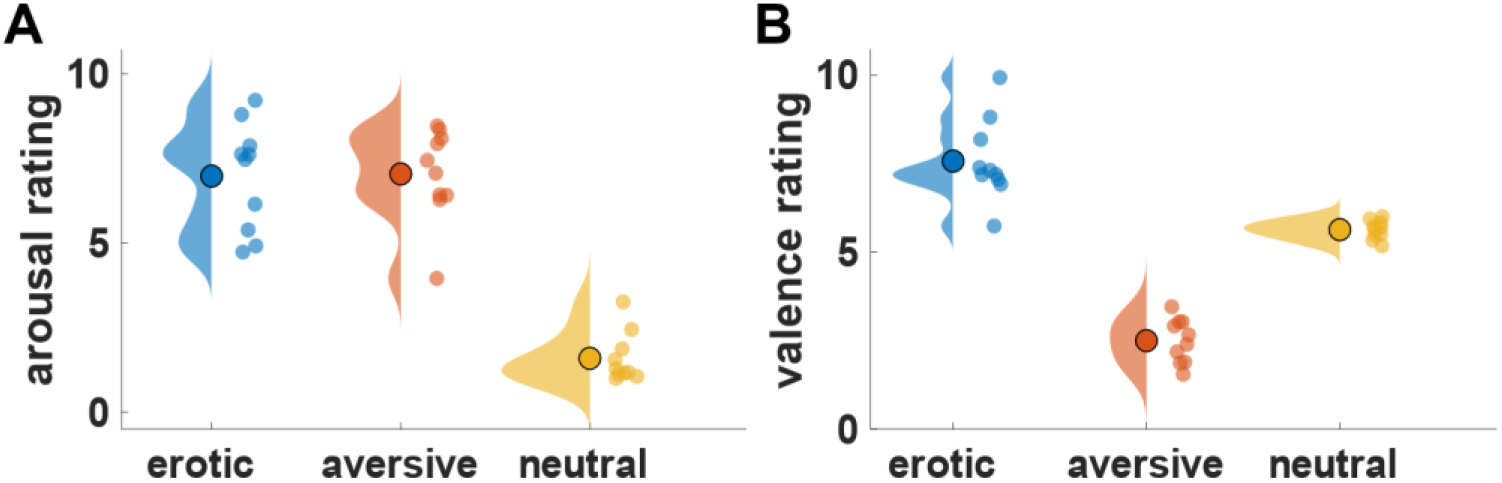
Pilot study image ratings (n = 10) of erotic, aversive and neutral cues (**A:** Arousal; **B:** Valence); Small colored dots depict single-subject means.

### Physical properties of stimulus material used in the study

Using MATLABS SHINE toolbox, images of all three experimental conditions were matched with respect to mean pixel intensity (*F*[2, 285] = 2.03, *p* = .133; mean ± SD = erotic: 0.42 ± 0.001; neutral: 0.42 ± 0.001; aversive: 0.42 ± 0.001) and pixel contrast (*F*[2,285] = 1.06, *p* = .347; mean ± SD = erotic: 0.19 ± 0.012; neutral: 0.19 ± 0.001; aversive: 0.19 ± 0.01) (Figure S2).

**Supplementary Figure S2.**
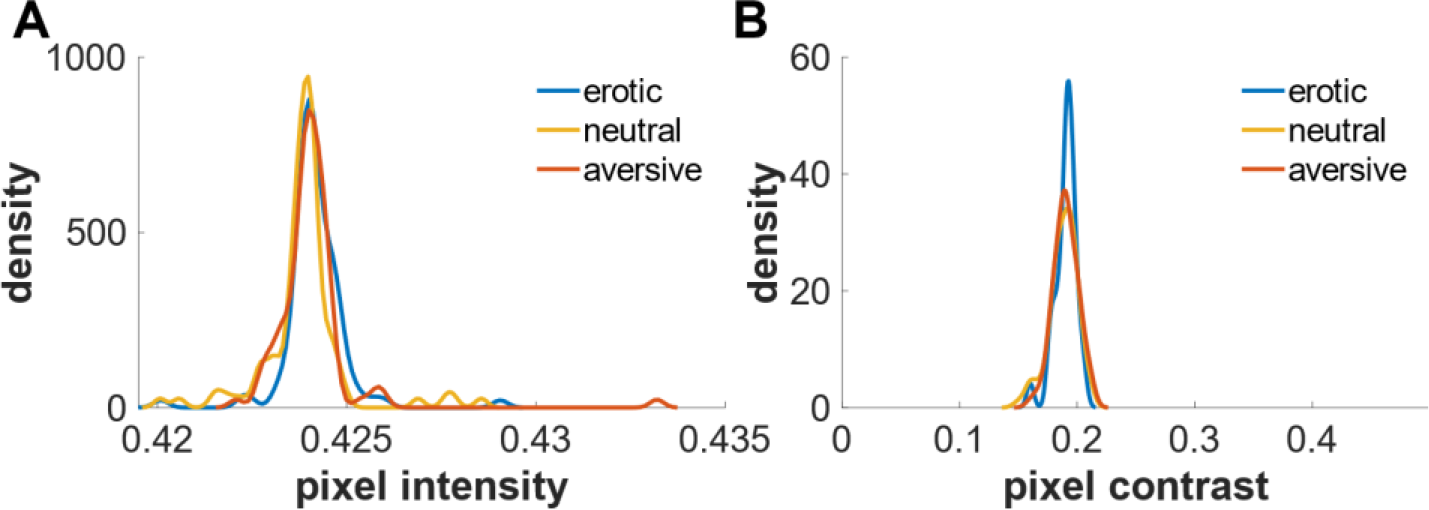
Physical properties of erotic, aversive and neutral cues (**A**: Mean pixel intensity; **B**: Standard deviation of pixel intensities (contrast)).

**Supplementary Table S1.** Pairwise comparisons of cue-evoked mean pupil dilation (Interaction effect: time bin (0.5s/bin) x condition).

| Time Bin | Condition (1) | Condition (2) | Difference | StdErr | *p-value* | Lower bound | Upper bound |
| --- | --- | --- | --- | --- | --- | --- | --- |
| 1 | erotic | aversive | 0.002 | 0.0001 | .287 | 0.001 | 0.004 |
| 1 | erotic | neutral | 0.002 | 0.001 | .298 | 0.001 | 0.004 |
| 1 | aversive | neutral | 0.0002 | 0.001 | 1.000 | -0.003 | 0.003 |
| 2 | erotic | aversive | 0.02 | 0.002 | < .001* | 0.013 | 0.024 |
| 2 | erotic | neutral | 0.03 | 0.002 | < .001* | 0.023 | 0.035 |
| 2 | aversive | neutral | 0.01 | 0.002 | < .001* | 0.007 | 0.015 |
| 3 | erotic | aversive | 0.04 | 0.004 | < .001* | 0.027 | 0.045 |
| 3 | erotic | neutral | 0.05 | 0.004 | < .001* | 0.037 | 0.058 |
| 3 | aversive | neutral | 0.01 | 0.003 | < .001* | 0.005 | 0.018 |
| 4 | erotic | aversive | 0.05 | 0.005 | < .001* | 0.036 | 0.059 |
| 4 | erotic | neutral | 0.06 | 0.005 | < .001* | 0.047 | 0.072 |
| 4 | aversive | neutral | 0.01 | 0.002 | < .001* | 0.006 | 0.018 |
| 5 | erotic | aversive | 0.05 | 0.005 | < .001* | 0.038 | 0.062 |
| 5 | erotic | neutral | 0.06 | 0.006 | < .001* | 0.046 | 0.074 |
| 5 | aversive | neutral | 0.01 | 0.002 | < .001* | 0.005 | 0.016 |
| 6 | erotic | aversive | 0.05 | 0.005 | < .001* | 0.033 | 0.059 |
| 6 | erotic | neutral | 0.06 | 0.006 | < .001* | 0.042 | 0.070 |
| 6 | aversive | neutral | 0.01 | 0.002 | < .001* | 0.004 | 0.017 |
| 7 | erotic | aversive | 0.04 | 0.005 | < .001* | 0.027 | 0.052 |
| 7 | erotic | neutral | 0.05 | 0.006 | < .001* | 0.034 | 0.062 |
| 7 | aversive | neutral | 0.01 | 0.002 | .002* | 0.003 | 0.014 |
| 8 | erotic | aversive | 0.04 | 0.005 | < .001* | 0.023 | 0.047 |
| 8 | erotic | neutral | 0.04 | 0.005 | < .001* | 0.030 | 0.056 |
| 8 | aversive | neutral | 0.01 | 0.002 | .005* | 0.002 | 0.014 |
| 9 | erotic | aversive | 0.03 | 0.004 | < .001* | 0.020 | 0.042 |
| 9 | erotic | neutral | 0.04 | 0.005 | < .001* | 0.025 | 0.050 |
| 9 | aversive | neutral | 0.01 | 0.003 | .054 | 0.000 | 0.013 |
| 10 | erotic | aversive | 0.03 | 0.004 | < .001* | 0.019 | 0.040 |
| 10 | erotic | neutral | 0.04 | 0.004 | < .001* | 0.024 | 0.046 |
| 10 | aversive | neutral | 0.01 | 0.003 | .115 | 0.000 | 0.012 |

*Note.* Asterisks indicate significant correlations on the $p < .05$ level (Bonferroni corrected). StdErr = standard error; Lower and upper bounds describe the 95% confidence interval.

### Exploratory analysis on skin conductance responses following image presentation

A repeated measures ANOVA indicated that while the sum of SCR amplitudes and mean phasic activity did not differ between experimental conditions (erotic, aversive, neutral), there was a higher mean number of evoked SCR’s following erotic images (mean ± Std = 83.31 ± 33.13) compared to aversive (mean ± Std = 77.83 ± 28.96) and neutral ones (mean ± Std = 78.06 ± 31.10) (Figure S3). However, none of the pairwise comparisons survived Bonferroni correction.

**Supplementary Figure S3.**
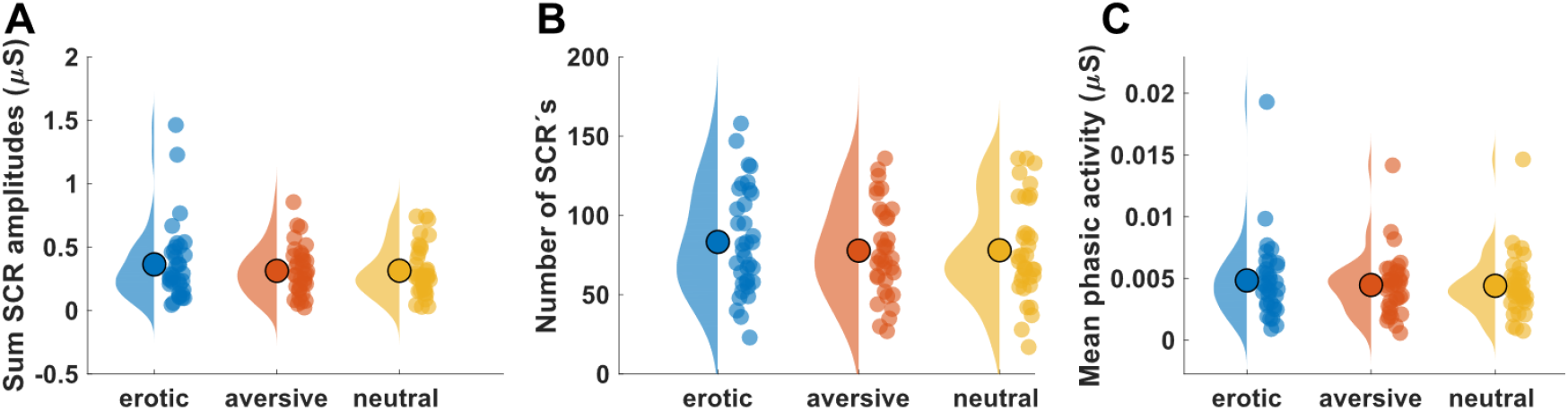
Sum of SCR amplitudes **(A)**, number of significant SCR’s **(B)** and mean phasic activity **(C)** in the interval 1-6 seconds post image onset; Small colored dots depict single-subject means; *µs* = microsiemens.

### Within-subject correlation of physiological measures in response to cue presentation

Within-subject cue-evoked changes in all three physiological measures (pupil size, heart rate, EDA) were at most weakly correlated (Figure S4, Table S2). Pupil size showed a small but significant mean correlation with heart rate change in response to image presentation (*r*_mean_ = .09, *p* < .001). Single-subject data indicated that while some subjects showed substantial associations, correlations were around zero in other participants. In contrast, pupil- and heart rate change both showed no associations with phasic electrodermal activity following image onset (Pupil / EDA: *r*_mean_ = .02, *p* = .142; HR / EDA: *r*_mean_ = -.03, *p* = .227).

**Supplementary Figure S4.**
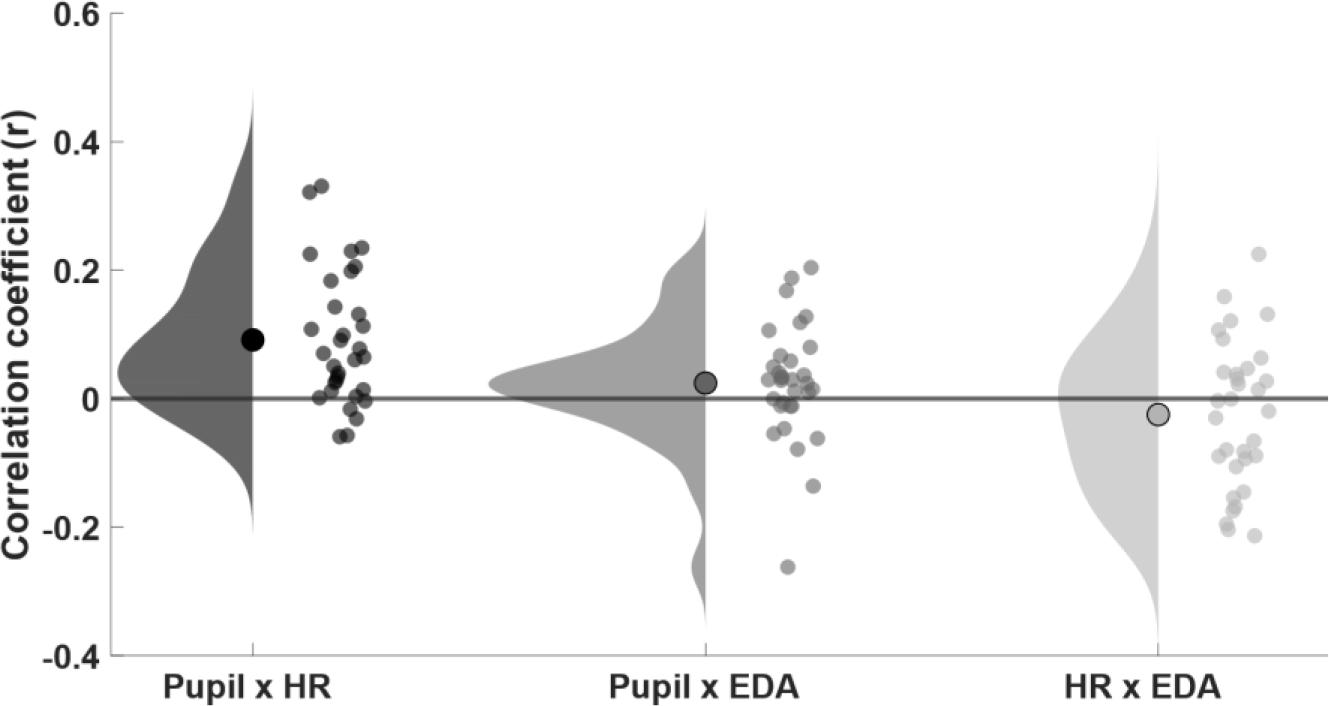
Mean within-subject associations (*r*) between physiological measures (pupil, heart rate (HR), electrodermal activity (EDA)) in response to cue presentation; Small gray dots depict single-subject coefficients.

**Supplementary Table S2.** Correlation statistics (rmeans) quantifying single-trial concordance between physiological measures (pupil, heart rate (HR), electrodermal activity (EDA)).

| Measures | $r_{\text{mean}}$ | CI | $p$-value |
| --- | --- | --- | --- |
| **Pupil x HR** | .09 | [0.05; 0.13] | < .001* |
| **Pupil x EDA** | .02 | [-0.01; 0.06] | .142 |
| **HR x EDA** | -.03 | [-0.07; 0.02] | .227 |

Note. Asterisks indicate significant correlations on the $p < .05$ level; $r$ = Pearson's correlation coefficient; CI = 95% confidence interval.

### Associations amongst physiological cue-reactivity indices

After we examined the trialwise associations between physiological measures in general, we assessed subjectś cue-reactivity response in every physiological measure as well as their concordance. To this end, we calculated difference scores between the mean response to erotic and neutral and between aversive and neutral content respectively. Next, we used Pearson’s correlations to quantify the association between those difference scores. As evident from Figure S5 (**A & D**), the above-mentioned selective increase in pupil diameter following erotic and aversive image content compared to neutral showed only small and non-significant associations to cardiovascular cue-reactivity. Subjects showing most pronounced differences in heart rate change between conditions exhibited only small cue-reactivity effects in pupil diameter (*r* = .26, *p* = .148). In addition, heightened cue-related pupil responses to aversive images were slightly and non-significantly associated with increased phasic electrodermal activity (Figure S5 (**E**); *r* = .19, *p* = .310). Detailed correlation statistics between physiological cue-response measures are depicted in Table S3.

**Supplementary Figure S5.**
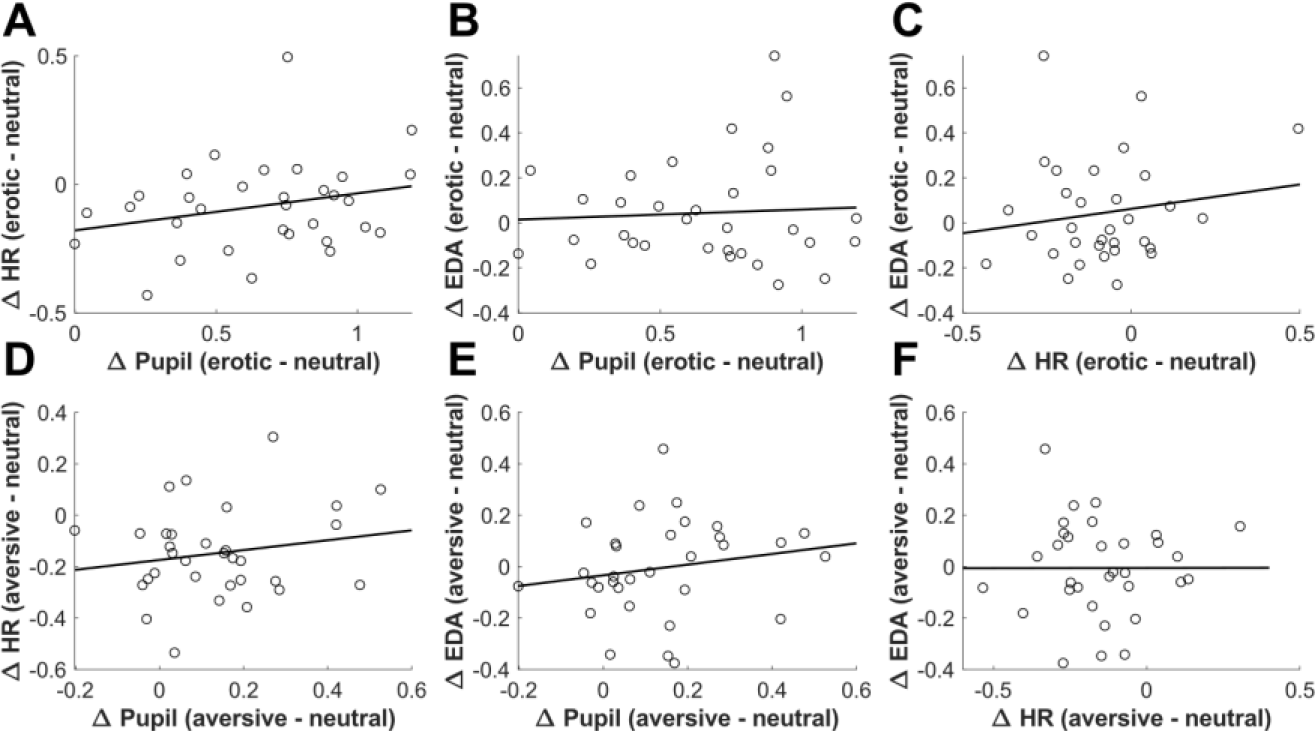
Associations amongst physiological cue-reactivity indices (difference scores; Pupil, Heart Rate (HR), Electrodermal Activity (EDA)).

**Supplementary Table S3.** Correlation statistics quantifying concordance between physiological cue-reactivity indices (difference scores; Pupil, Heart Rate (HR) & Electrodermal Activity (EDA)).

| Physiological measures | $\Delta$ Contrast | Correlation coefficient ($r$) | CI | $p$-value |
| --- | --- | --- | --- | --- |
| Pupil x HR | Erotic - neutral | .26 | [-0.10; 0.56] | .148 |
|  | Aversive - neutral | .18 | [-0.18; 0.50] | .315 |
| Pupil x EDA | Erotic - neutral | .06 | [-0.29; 0.40] | .740 |
|  | Aversive - neutral | .19 | [-0.17; 0.50] | .310 |
| HR x EDA | Erotic - neutral | .16 | [-0.20; 0.48] | .375 |
|  | Aversive - neutral | .002 | [-0.35; 0.35] | .993 |

Note. $r$ = Pearson's correlation coefficient; CI = 95% confidence interval.

### Associations between model-free and model-based measures of temporal discounting

**Supplementary Figure S6.**
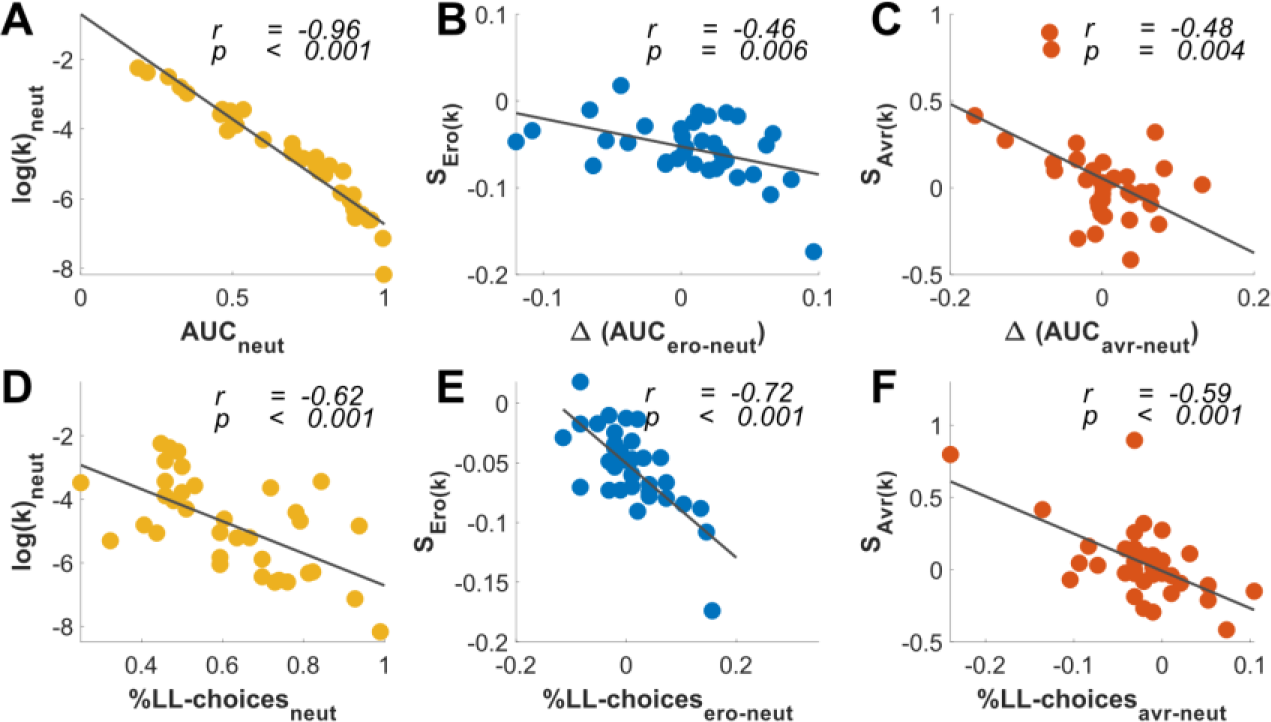
Associations between model-free (AUC, LL-choice proportions) and model-based measures (S_Ero(k)_, S_Avr(k)_) of discounting behavior. *r* = Pearson’s correlation coefficient.

**Supplementary Table S4.** Correlation statistics quantifying associations between model-free (AUC, LL-choice proportions) and model-based measures (S_Ero(k)_, S_Avr(k)_) of discounting behavior.

| Measure (1) | Measure (2) | Correlation coefficient ($r$) | CI | $p$-value |
| --- | --- | --- | --- | --- |
| $\text{Log}(k)_{\text{neut}}$ | $\text{AUC}_{\text{neut}}$ | -.96 | [-.98; -.93] | < .001* |
| | % LL-choices$_{S_{\text{neut}}}$ | -.62 | [-.79; -.36] | < .001* |
| $S_{Ero(k)}$ | $\Delta (\text{AUC}_{\text{ero-neut}})$ | -.46 | [-.69; -.15] | .006 |
| | % LL-choices$_{S_{\text{ero-neut}}}$ | -.72 | [-.85; -.51] | < .001* |
| $S_{Avr(k)}$ | $\Delta (\text{AUC}_{\text{avr-neut}})$ | -.48 | [-.70; .17] | .004 |
| | % LL-choices$_{S_{\text{avr-neut}}}$ | -.59 | [-.77; -.32] | < .001* |

Note. Asterisks indicate significant correlations on the $p < .05$ level; $r$ = Pearson's correlation coefficient; CI = 95% confidence interval.

### Association of behavioral (shift-parameters) and pupillary cue-reactivity indices (difference scores)

The magnitude of erotic shift-parameter on log(*k*) (*S_Ero(k)_*) showed a slightly non-significant negative association with differential pupil responses to erotic stimuli, indicating that subjects exhibiting increased responses to erotic images also showed a more pronounced decrease in their discounting behavior in the erotic condition (*r* = -.29, *p* = .096; Figure S7, **A**). There was no comparable association for the aversive condition (*r* = .11, *p* = .546). In addition, elevated pupil responses to affective stimulus material (erotic & aversive) were positively associated with respective *ß*-shift parameters (*S_Ero(ß)_, S_Avr(ß)_*; Figure S7, **B** & **D**) assuming that an increased physiological cue-response was associated with less decision noise. Despite non-significant, this effect was more pronounced for aversive image condition (Aversive: *r* = .22, *p* = .220; Erotic: *r* = .13, *p* = .487). Detailed correlation statistics are depicted in Table S5.

**Supplementary Figure S7.**
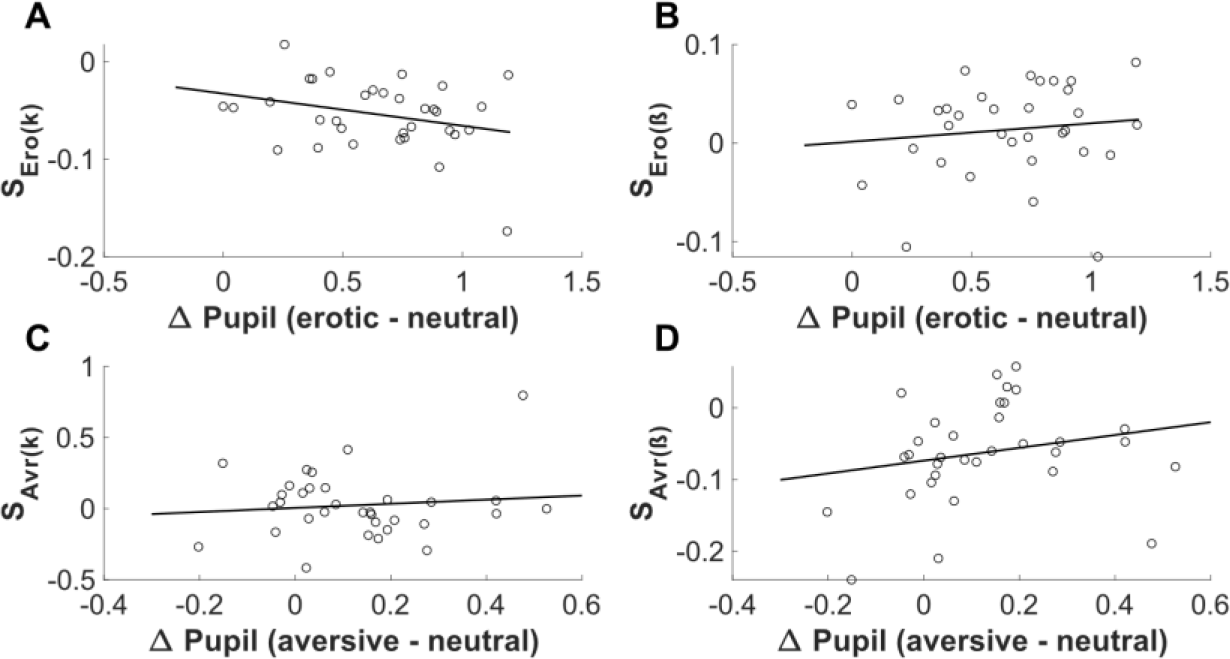
Association between pupillary cue-reactivity indices (difference scores) and erotic- (**A, B**) and aversive (**C, D**) shift-parameters (*S_Ero(k,ß)_*, *S_Avr(k,ß)_*).

**Supplementary Table S5.** Correlation statistics quantifying the association between behavioral (shift-parameters; (*S_Ero(k,ß)_*, *S_Avr(k,ß)_*)) and pupillary cue-reactivity indices (difference scores).

| Measures | Correlation coefficient ($r$) | CI | $p$-value |
| --- | --- | --- | --- |
| $\Delta \text{Pupil}_{(\text{ero-neut})} \times S_{\text{Ero}(k)}$ | -.29 | [-0.58; 0.05] | .096 |
| $\Delta \text{Pupil}_{(\text{avr-neut})} \times S_{\text{Avr}(k)}$ | .11 | [-0.24; 0.44] | .546 |
| $\Delta \text{Pupil}_{(\text{ero-neut})} \times S_{\text{Ero}(\beta)}$ | .13 | [-0.23; 0.45] | .487 |
| $\Delta \text{Pupil}_{(\text{avr-neut})} \times S_{\text{Avr}(\beta)}$ | .22 | [-0.13; 0.52] | .220 |

Note. $r$ = Pearson's correlation coefficient; CI = 95% confidence interval.

### Habituation of physiological cue response

We assessed habituation processes of evoked physiological responses to image presentation over the course of the experiment. To this end, we separately z-scored and binned pupil, heart rate and electrodermal activity data (trial means) into three trial bins. Using separate repeated measures ANOVA, we compared evoked physiological responses over the course of the experiment (within factor: time bin). As shown in Figure S8 (**A**), evoked pupil responses slightly decreased over trial bins (F[1, 32] = 4.18, *p* = .028, *η*^2^= 0.12, *ε* = 0.82). Similar patterns were numerically apparent for evoked heart rate responses and phasic electrodermal activity data (EDA) but both did not reach significance (Heart Rate: F[1, 33] = 1.46, *p* = .240; **B;** EDA: F[1, 34] = 0.58, *p* = .498, *ε* = 0.66; **C**).

**Supplementary Figure S8.**
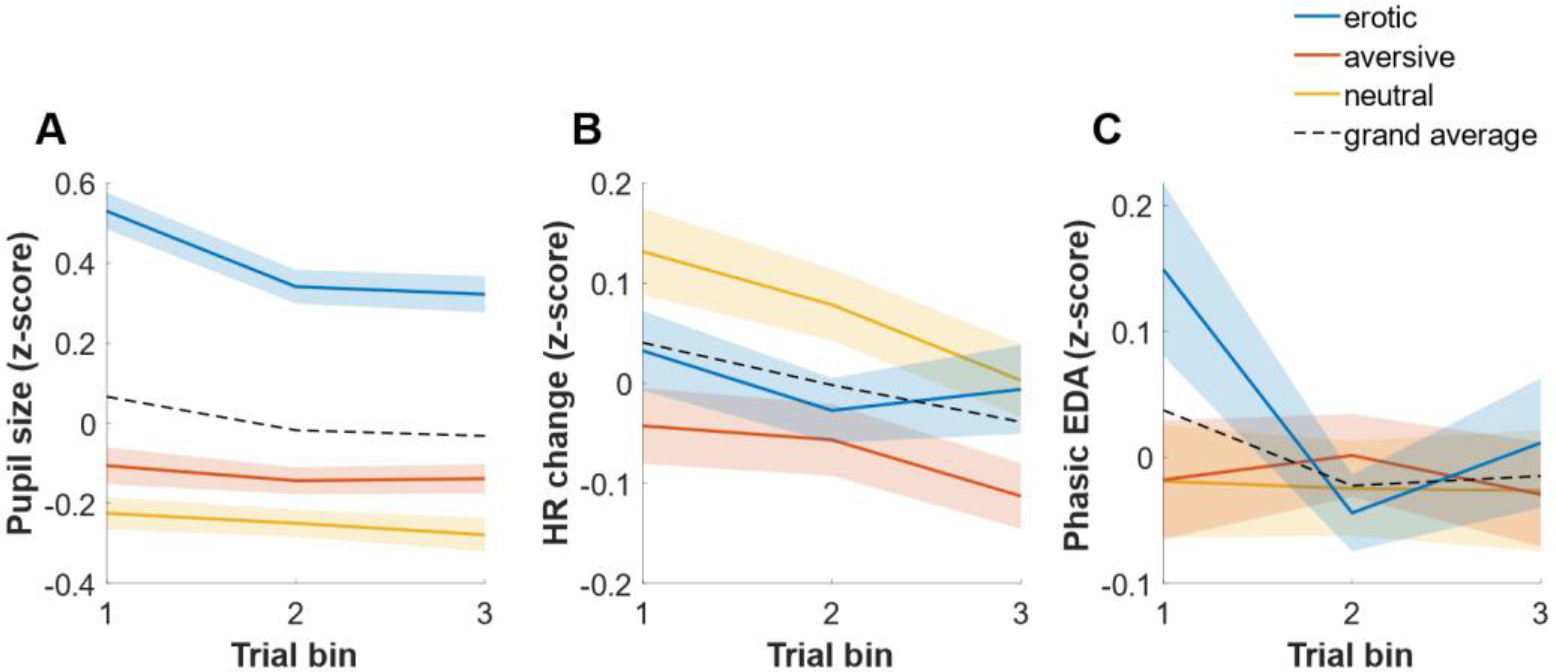
Cue-evoked physiological responses over the course of the experiment; Valid trials of pupil size (**A**), heart rate (HR; **B**) and phasic electrodermal activity data (EDA; **C**), were binned into three bins separately for neutral, erotic and aversive cue conditions; Shaded areas depict standard errors (SE).

### Evaluation of physiological and behavioral indices of cognitive effort

#### Analysis

As pupil dilation appears sensitive to various internal states or cognitive manipulations, including mental arithmetic, memory processes and decision formation (van der Wel & van Steenbergen, 2018), we hypothesized, that high decision conflict during temporal discounting would likewise result in an elevated pupil dilation indicating high cognitive effort irrespective of cue condition. We examined this using three complementary approaches. First, mean pupil size starting at larger later reward (LL) onset until decision screen onset was compared between easy, medium and hard trials using repeated measures ANOVA. For every subject we determined trial difficulty by classifying all trials into three terciles depending on the numeric subjective value (SV) difference between smaller sooner and the larger later options. Second, as cognitive conflict might cumulate and reach its peak immediately before a decision between SS and LL-options is made, we used a linear mixed model to analyze whether median pupil size in the interval 1 second prior to decision (button press) differed between trial difficulty levels (easy, medium, hard). We added cue condition (erotic, aversive, neutral) as well as the interaction of cue condition and trial difficulty as fixed effects while fitting a random effect for subject. Third, using a Pupil Response Estimation Toolbox (PRET) we constructed a general linear model (GLM) to examine trial difficulty- and cue-effects on pupil dilation (Denison et al., 2020). We defined two sustained boxcar regressors, spanning the periods of image- and LL-reward presentation. Both regressors were convolved with a pupil response function (PRF) of the form:

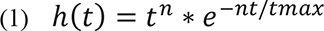

Here, *h* is the predicted pupil size, *t* is the time in ms, *n* controls the shape of the function, and *tmax* controls its temporal scale as it is the time of the maximum (Hoeks & Levelt, 1993). To examine interaction effects, single trial data were fitted separately for all cue conditions and trial difficulty levels. Fitted boxcar amplitudes for LL-reward presentation were compared using rmANOVA (within factors: cue condition, trial difficulty).

As trial difficulty might also affect behavioral markers sensitive to cognitive effort, like reaction times (RT), we again used the above mentioned linear mixed model to assess effects of trial difficulty, experimental condition and their interaction (fixed effects) on reaction times, fitting a random effect for every subject. To investigate whether trial difficulty effects on behavioral and physiological indices are associated on the subject level, we extracted single-subject fixed effect estimates from the above mentioned linear mixed models on pupil and reaction times and implemented a Pearson’s correlation analysis.

## Results

Pupil trajectories in response to easy, medium and hard trials are depicted in Figure S9 (**A**). There was a substantial pupil dilation in response to larger later reward onset which reached a plateau approximately 1.5 seconds post LL presentation. Repeated measures ANOVA revealed that trial difficulty did not further change mean evoked pupil response (F[2,64] = 0.12, *p* = .830, *ε* = 0.76).

**Supplementary Figure S9.**
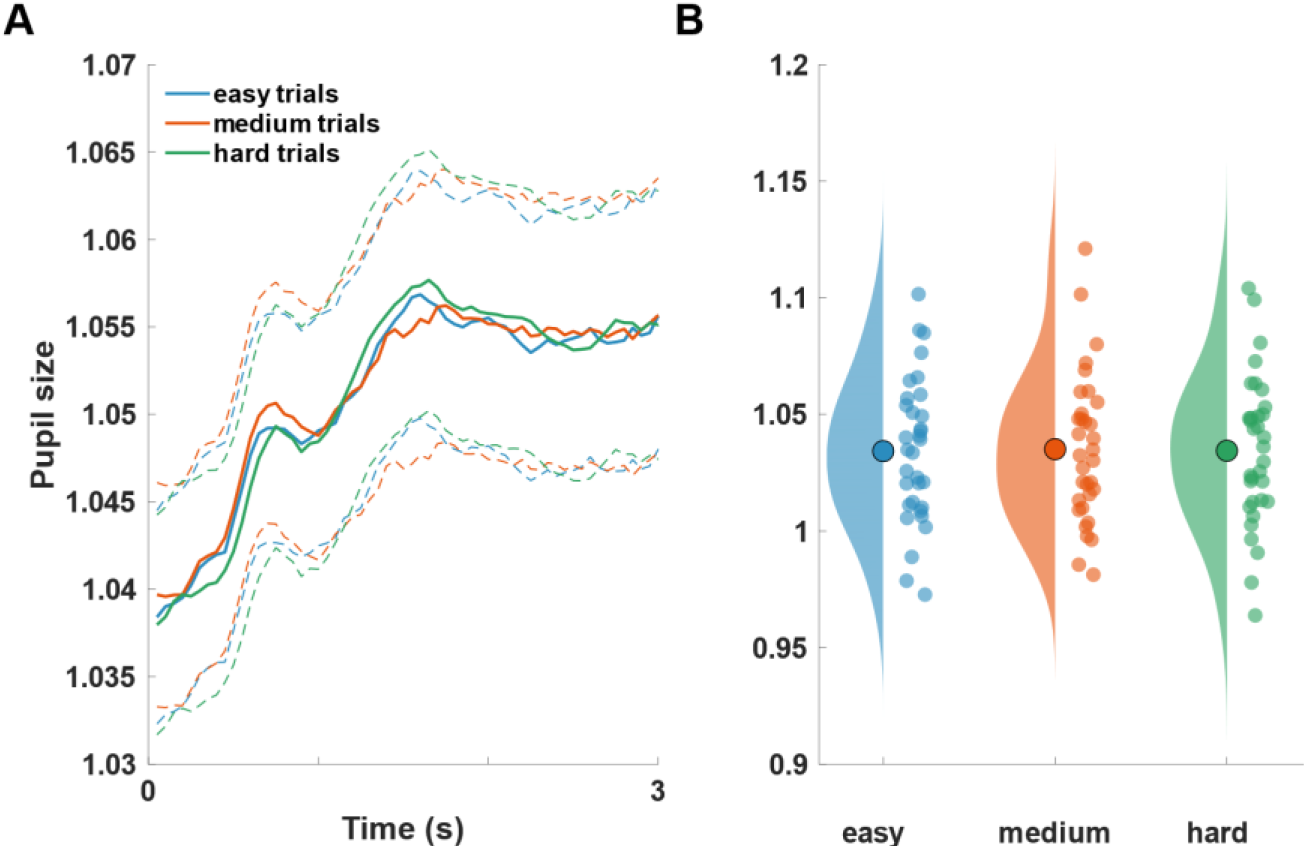
(A) Grand average pupil trajectories as a function of trial difficulty following larger later (LL) reward onset; dotted lines depict standard errors (SÉs) **(B)** Mean single-subject pupil responses for the interval (LL-onset until decision screen onset).

To assess mental effort effects on pupil dilation in more detail, we used a linear mixed model to quantify trial difficulty and condition effects on median pupil size within the last second interval prior to decision. Complementing previous results, we identified a significant main effect for experimental condition F[1,261], *p* < .001, replicating enhanced pupil responses following erotic and aversive images compared to neutral. There was no further significant main effect for trial difficulty (F[1,261], *p* = .375) or interaction effect between trial difficulty and condition (F[1,261], *p* = .451) in that specific trial period. Two sustained boxcar regressors spanning the intervals of image- and LL-reward presentation, each convolved with a pupil response function, were used to model trialwise pupil trajectories. As trial difficulty effects should first emerge following LL-presentation, we compared boxcar amplitudes for this second regressor using repeated measures ANOVA (within factors: Cue condition, Trial difficulty). While results from the general linear model (GLM) confirmed the missing trial difficulty effect on pupil size (F[2,64] = 0.09, *p* = .922) we found a significant main effect of cue condition indicating that this second dilatory process was most pronounced for neutral image condition, followed by aversive and erotic, possibly indicating a ceiling effect (F[2,64] = 21.63, *p* < .001; mean ± SD: neutral = 2.05 ± 2.01, aversive = 1.67 ± 1.94, erotic = 0.53 ± 0.78). Raw and modeled pupil trajectories as well as boxcar amplitudes split by difficulty levels are depicted in Figure S10.

**Supplementary Figure S10.**
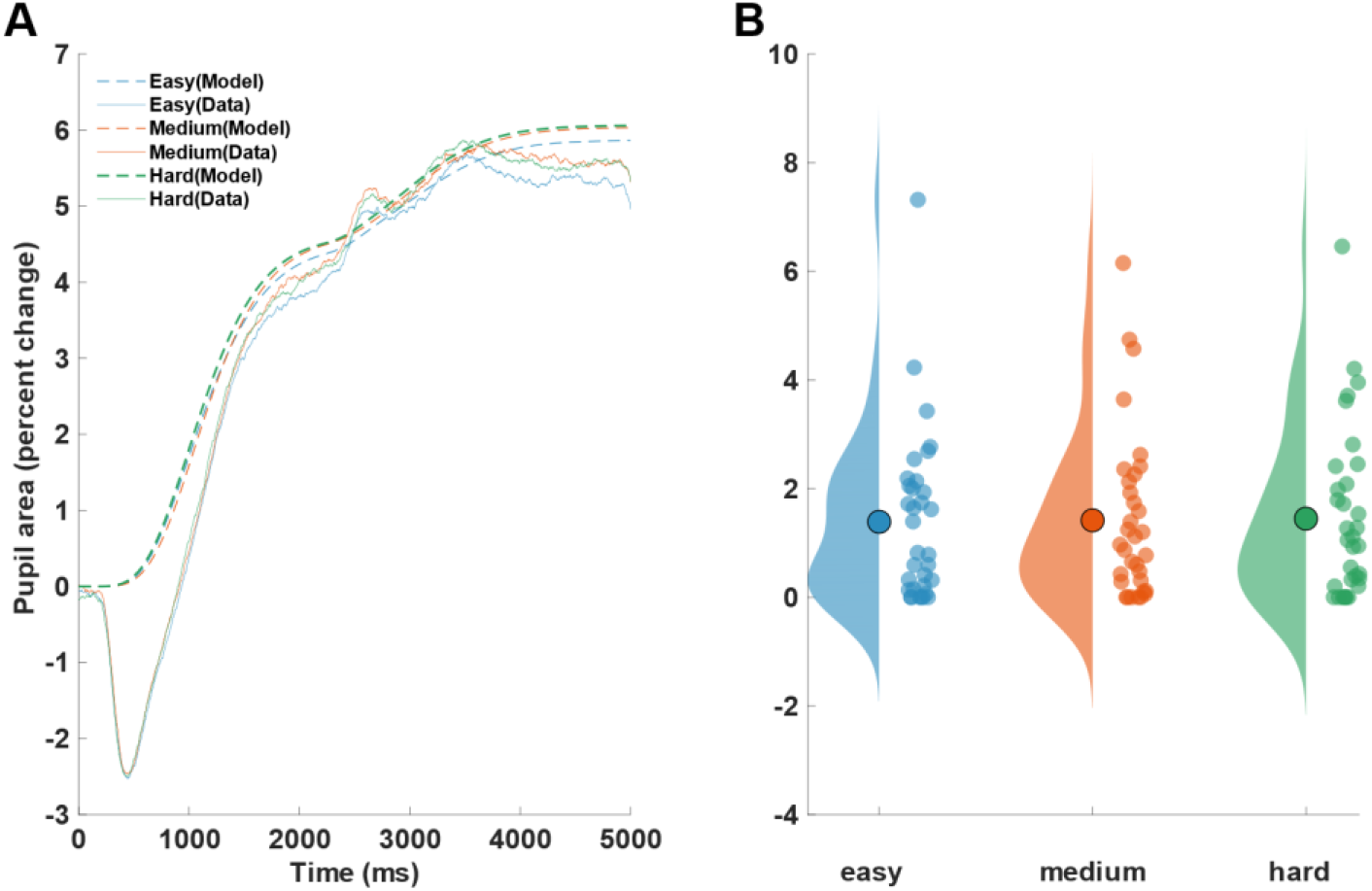
(A) Grand average pupil trajectories as a function of trial difficulty following image onset; solid/dotted lines = real/modeled data (GLM); **(B)** Unstandardized regression weights of LL-reward presentation on pupil size split by trial difficulty level.

As cognitive effort was not visible in differential physiological responses (pupil dilation) we explored whether trial difficulty was captured by behavioral markers of cognitive effort. To this end, we again used a linear mixed model to assess effects of trial difficulty and experimental condition (fixed effects) on reaction times (RT). Results showed a significant main effect of trial difficulty level on reaction times (F[1,277], *p* = .029), indicating slower RT’s for medium and hard trials compared to neutral (medium vs easy: *t*(6.514), *p* < .001); hard vs easy: *t*(6.08), *p* < .001, see Figure S11). RT’s for hard and medium trials did not differ significantly (*t*(−0.43), *p* = .902). There were no further significant main or interaction effects.

**Supplementary Figure S11.**
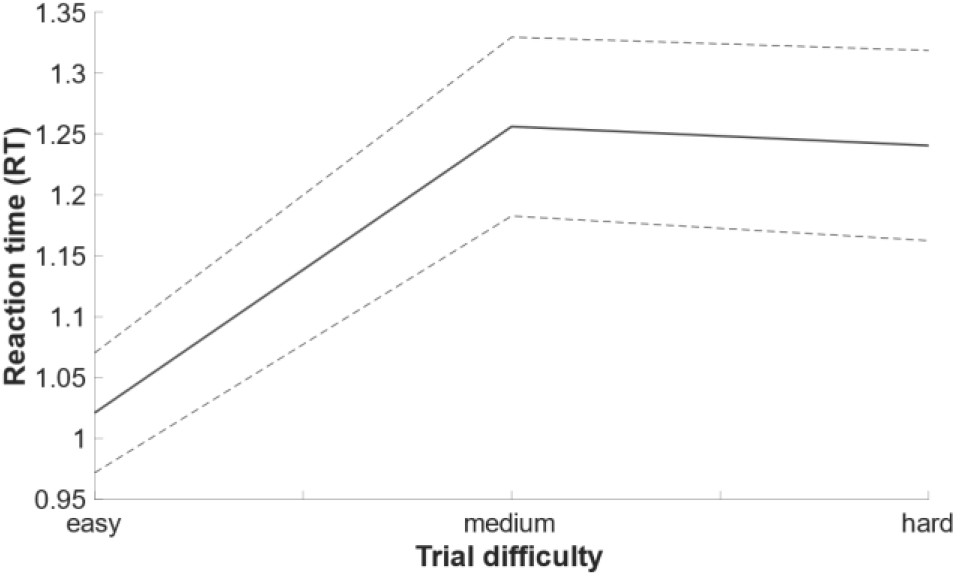
Reaction time (RT) differences as a function of trial difficulty (easy, medium, hard). Dotted lines depict grand mean across all participants ± standard errors (SE’s).

We also tested whether behavioral (RTs) and physiological (pupil dilation) indices of cognitive effort showed associations on the subject level. By this means, we extracted single-subject fixed effect estimates from the above mentioned linear mixed models on pupil and reaction times and implemented a Pearson’s correlation analysis. As shown in Figure S12, there seemed to be no strong interdependency between both measures (*r* = -.20, *p* = .260, CI _(95%)_ = [-.51; .15]).

**Supplementary Figure S12.**
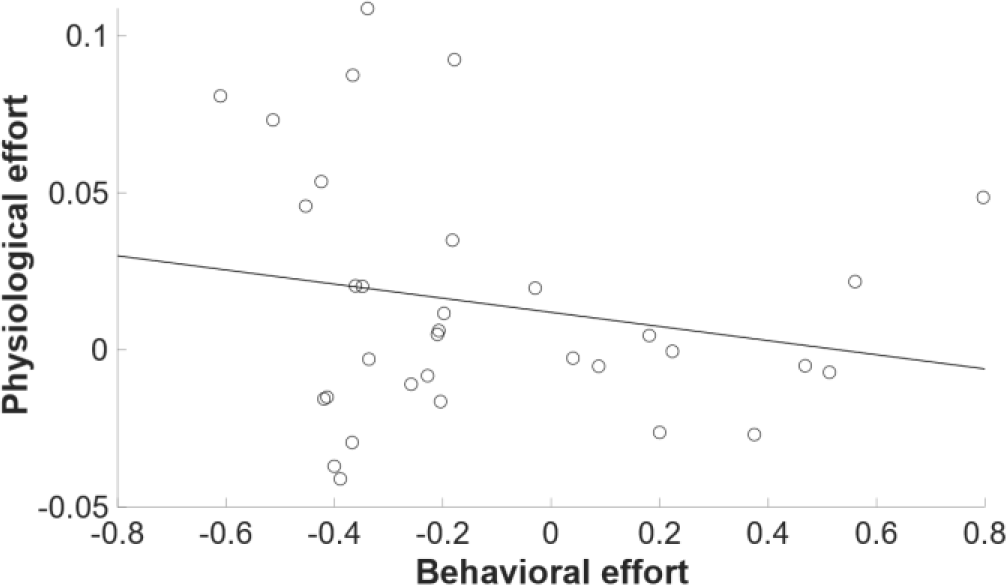
Physiological (pupil size) and behavioral (RT) indices of cognitive effort and their association. Dots depict single subject beta coefficients from the mixed models: **(1)** RT ∼ difficulty + condition + difficulty*condition + (1|Subject); **(2)** Pupil Size ∼ difficulty + condition + difficulty*condition + (1|Subject).

### Association of working memory capacity (WMC) and temporal discounting

As working memory capacity (WMC) is often negatively associated with measures of choice impulsivity like temporal discounting we tested whether this is the case in our data. We calculated a working memory compound score for every participant, that is the mean z-score from four different working memory tasks (forward/backward digit span, operation/listening span; Redick et al., 2012; van den Noort et al., 2008; Wechsler, 2008). Next, we implemented Pearson’s correlation between working memory scores and estimated neutral log(*k*)-parameters from the computational shift model (see Eqs. 4 & 5). Results showed no substantial association between WMC and estimated neutral log(*k*)-parameters (*r* = -.11, *p* = .526, CI _(95%)_ = [-.43; .23]; Figure S13).

**Supplementary Figure S13.**
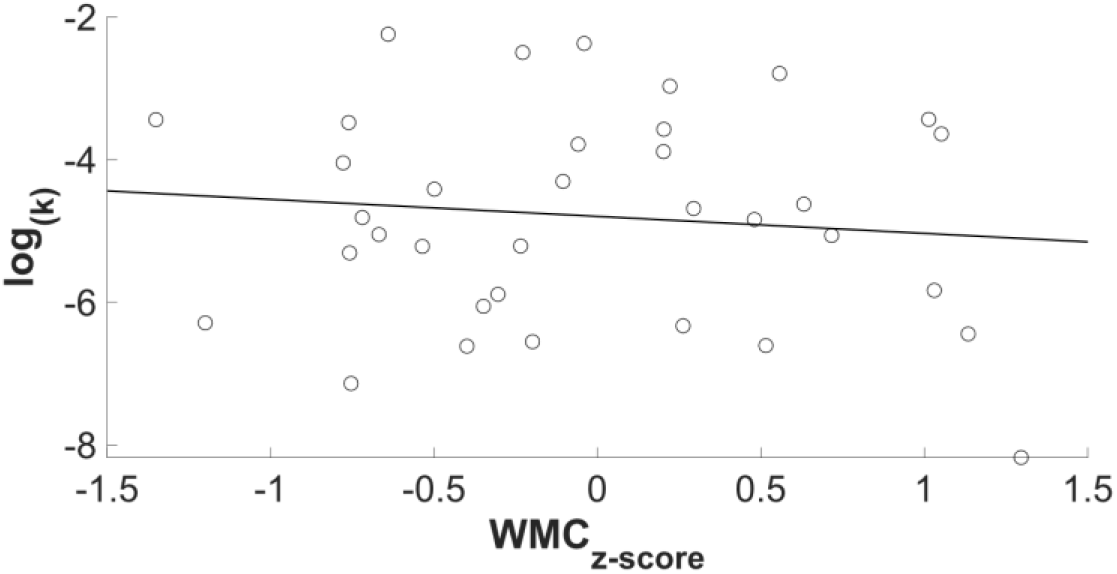
Association between mean standardized working memory score (WMC) and neutral log (*k*)-parameter from the computational shift-model.

